# A Thermodynamic Theory of Axon Guidance: Navigation Through High-Entropy Signaling States

**DOI:** 10.64898/2026.09.22.752727

**Authors:** William G. Wadsworth

## Abstract

Precise neural circuit formation requires growth cones to integrate multiple, sometimes competing, guidance signals into persistent yet adaptable movement. Here, we propose a theoretical framework that recasts axon guidance as a thermodynamically regulated computation. An artificial neural network, trained on in vivo genetic data and associated outgrowth patterns, maps the microstates of a localized guidance signaling network to macroscopic outgrowth behaviors. Using the trained model, we simulated axon pathfinding through extracellular gradients of molecular cues and created dynamic entropic landscapes. These results reveal that growth cones navigate high-entropy ridges that preserve plasticity. Navigation along these ridges supports persistent extension, whereas turning and branching occur near boundaries between competing entropic macrostates, consistent with transitions in the cytoskeletal machinery that controls growth-cone movement. More broadly, the framework proposes that robust biological patterning can emerge from microscopic variability when signaling networks operate near boundaries between competing behavioral states.

**Significance Statement:** How do complex biological systems remain robust yet adaptable? We address this fundamental question by conceptualizing axon guidance as a thermodynamic process. By embedding a neural-network predictive model of signal integration within a statistical physics framework, we show computationally that migrating axonal growth cones need not passively settle into stable states, but can instead navigate fluctuating, high-entropy states. This thermodynamic strategy reconciles persistence with plasticity, allowing directed movement while keeping growth cones poised near critical boundaries where small biochemical fluctuations can trigger rapid turning or branching. More broadly, this framework suggests a general principle by which complex cellular systems use network degeneracy and fluctuations to coordinate robust behavior with rapid adaptation during migration, pattern formation, and tissue development.

## Introduction

The proper functioning of a nervous system relies on the establishment of an intricate network of precise, stereotypical connections. The mechanisms by which axon growth cones navigate to create the patterns required to make the needed connections must be versatile enough to generate a diverse repertoire of morphological patterns. This structural plasticity is regulated by neuronal gene expression and the responses it mediates to extracellular factors. By regulating how mechanical forces are applied to the plasma membrane, this regulatory system causes navigational behaviors, such as branching and turning. It must also regulate the spatiotemporal sequence of these behaviors to create the required patterns of outgrowth.

The canonical paradigm of axon guidance postulates that growth cones act as sensor-integrators, navigating toward targets by responding to extracellular cue gradients and transducing attractant and repellent cue signals into cytoskeletal dynamics (1–3). Historically, this view treats neurons as recipients of a spatial ‘blueprint’, a molecular roadmap that growth cones sense and follow. However, this model struggles to capture the full spatiotemporal complexity of neural circuit assembly. In part, this is because of complexity. Growth cones must continuously integrate noisy, often conflicting, extracellular cues to execute macroscopic morphological decisions. Moreover, from a biophysical perspective, the growth cone’s cytoskeleton is not merely a passive recipient of signaling pathways but rather operates as active matter: a non-equilibrium, energy-consuming network. As active matter, the cytoskeleton can undergo rapid mechanical phase transitions, such as spontaneous symmetry breaking, in which sudden asymmetric reorganization of cytoskeletal filaments can redirect outgrowth force. From this perspective, understanding how pathfinding emerges requires conceptualizing the growth cone not simply as a gradient-tracking machine but as a complex dynamical system.

In systems biology and physics, attractor-based models are used to analyze and predict the long-term behavior, stability, and state transitions of complex, non-linear systems (4–10). Traditionally, an attractor represents a highly robust, preferred macroscopic state. In free-energy or Waddington landscapes, attractors correspond to basins of stability where the effective free energy is minimized, and the biological state is more likely to be maintained. While these landscapes are useful for describing the establishment of a static state, such as cell fate or stable morphology, they are less apt for describing situations that require dynamic plasticity, such as spatiotemporal morphological patterning during development.

Here, we conceptualize axon guidance not as a process of settling into stable attractors, but rather as one of actively avoiding them. The network analyzed in this study consists of highly conserved genes encoding extracellular ligands, transmembrane receptors, a cytoplasmic scaffolding regulator, and a transcription factor. Together, these components form the localized Guidance Signaling Network (GSN). From a statistical mechanics perspective, the ensemble of microstates corresponds to the complete set of molecular configurations that the GSN can assume. A molecular configuration includes expression levels, receptor distributions, and potential molecular interactions. The macrostates of the system are the observable physical outgrowth behaviors. Importantly, biological networks exhibit degeneracy, in which multiple distinct microstate configurations give rise to indistinguishable macrostates (11–15). Network degeneracy allows the system to maintain functional robustness despite underlying variability.

To ensure plasticity, biological systems with adaptable behaviors are often modeled as operating near critical phase transitions between ordered and disordered states (16–23). We propose that axon pathfinding emerges when the GSN evolves towards entropic phase boundaries. That is, rather than converging to the bottom of an energetic basin where entropy is minimized, axon pathfinding travels along the ridges between attractors. These boundaries are where the coarse-grained network attains maximal macrostate (Shannon) entropy. At these locations, the GSN reduces the effective free-energy barrier that constrains cytoskeletal active matter. This facilitates rapid cytoskeletal reorganization, which is necessary for changes in the direction of outgrowth. To test this, we developed a computational framework that translates in vivo genetic data into a localized entropic steering algorithm. A trained Multi-Layer Perceptron (MLP) Artificial Neural Network maps combinatorial genotypes of an 11-component GSN to 12 distinct morphological outgrowth patterns. The MLP then evaluates the volatility of the GSN across a biologically constrained perturbation ensemble of distinct network microstates. The results demonstrate that spatial and temporal pathfinding decisions strictly coincide with entropic phase transitions, and that complex, neural architectures can emerge from the degeneracy of the underlying signaling network.

## Results

### Genetic Regulation of Force Distribution

To begin to understand how genetic network states regulate the force-producing machinery of the neuron, the directional bias of axon outgrowth proximal to the HSN cell body was quantified (Figure 1A). The HSN neurons are born in the tail region of the *Caenorhabditis elegans* embryo and migrate anteriorly to the mid-body. During larval stages, a single axon extends to the ventral nerve cord. Mutations in guidance genes that alter the patterning of axon outgrowth affect the initial direction of axon outgrowth. We have published the percentage of outgrowth in the anterior, dorsal, posterior, and ventral directions for wild-type animals as well as for single, double, and triple loss-of-function mutants in known guidance genes (Figure 1B)(24–27). These genes are all known to play significant roles in determining axon outgrowth patterning (Figure 1C) (28). A set of four genes encodes extracellular/environmental cues secreted by cells distant from the HSN cell body. One of these genes is egl-20, a Wnt family member. *C. elegans* has multiple Wnt genes and Wnt receptors. The egl-20 Wnt is secreted posterior to the HSN neuron, while other Wnt genes are expressed as a series of partially overlapping domains along the anterior-posterior axis (29). These genes affect HSN axon outgrowth from sources anterior to the cell body. To develop a model for the collective properties of these genes, the proxy, “WL,” is used to designate the collective effect of these Wnt ligands on outgrowth, and “WR” for the collective impact of their receptors.

**Figure 1.**
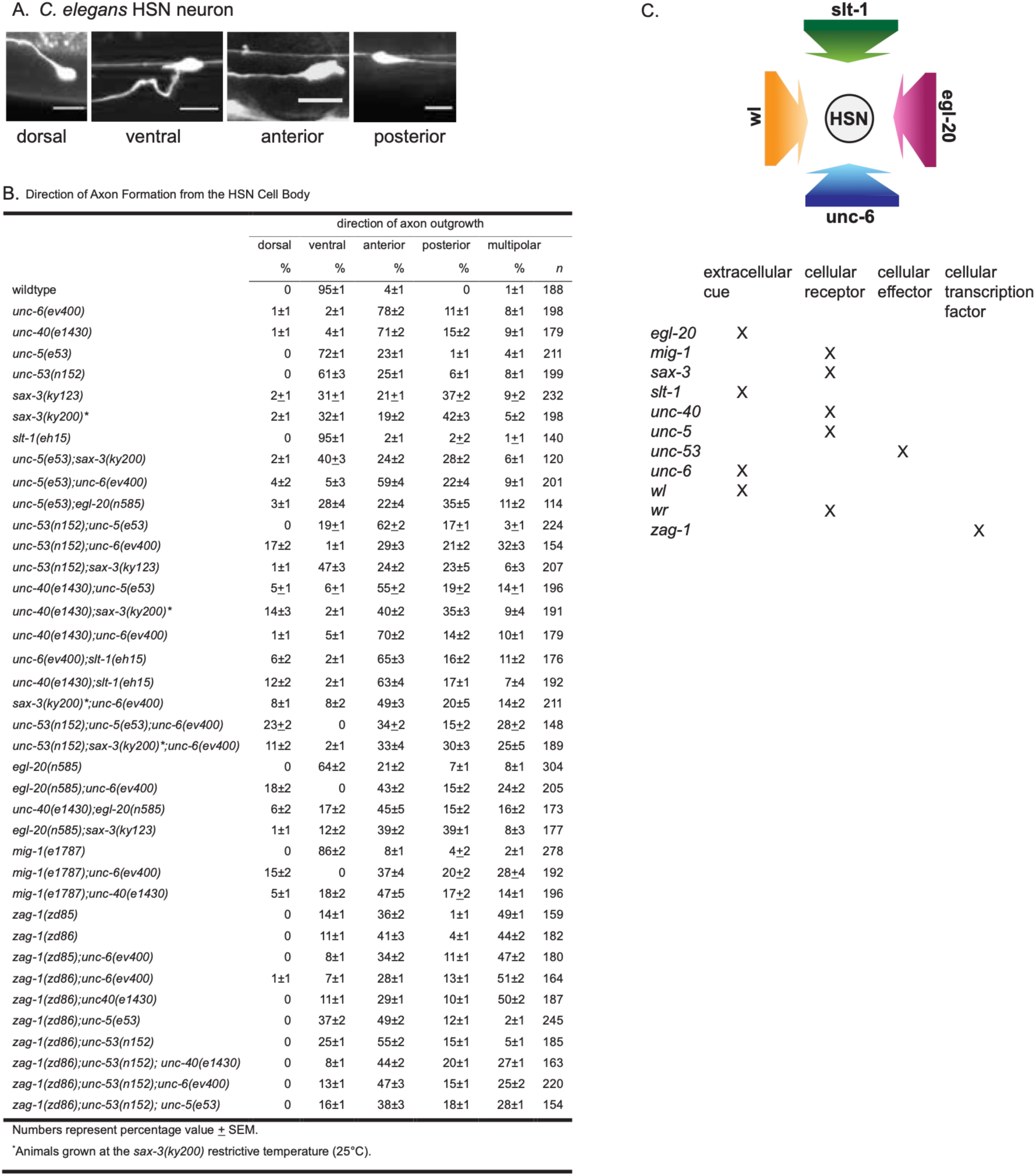
Genetic regulation and extracellular cues dictate the directional bias of HSN axon outgrowth. (A) Photomicrographs of L4 stage animals showing examples of axon outgrowth from the HSN cell body in wild-type and mutant animals. Ventral is down, and anterior is to the left. Scale bar: 10 µm. In wild-type animals, the HSN axon protrudes ventrally from the cell body. In mutants, the HSN axon can protrude in the anterior, posterior, or dorsal directions. Reproduced from Tang X, Wadsworth WG (2014) SAX-3 (Robo) and UNC-40 (DCC) Regulate a Directional Bias for Axon Guidance in Response to Multiple Extracellular Cues. PLoS ONE 9(10): e110031. https://doi.org/10.1371/journal.pone.0110031. (B) Table of HSN axon outgrowth, quantifying the percentage of initial outgrowth in the anterior, dorsal, posterior, and ventral directions for wild-type animals as well as single, double, and triple loss-of-function mutants in known guidance genes as reported in the cited references. (C) Top: Diagram showing the location, relative to the position of the HSN neuron during axon outgrowth, of the sources that secrete the indicated extracellular cues. Arrows depict that gradients of the cues form in the environment where HSN outgrowth migration occurs. Bottom: Table detailing the functional roles of the products of the guidance genes used in this study.

To visualize how these fluctuations in force generation translate into directed movements, a biased random walk was used to model outgrowth (Figure 2A) (25). For each genotype, simulations on a lattice were performed in which the biologically derived probability distributions determined the direction of each step. To capture the morphological footprint of each genetic state, we superimposed 25 independent walks of 500 steps, all initiating from a common origin. These patterns serve as a readout of how the underlying genetic network governs the spatial probability of where physical force is applied to the plasma membrane.

**Figure 2.**
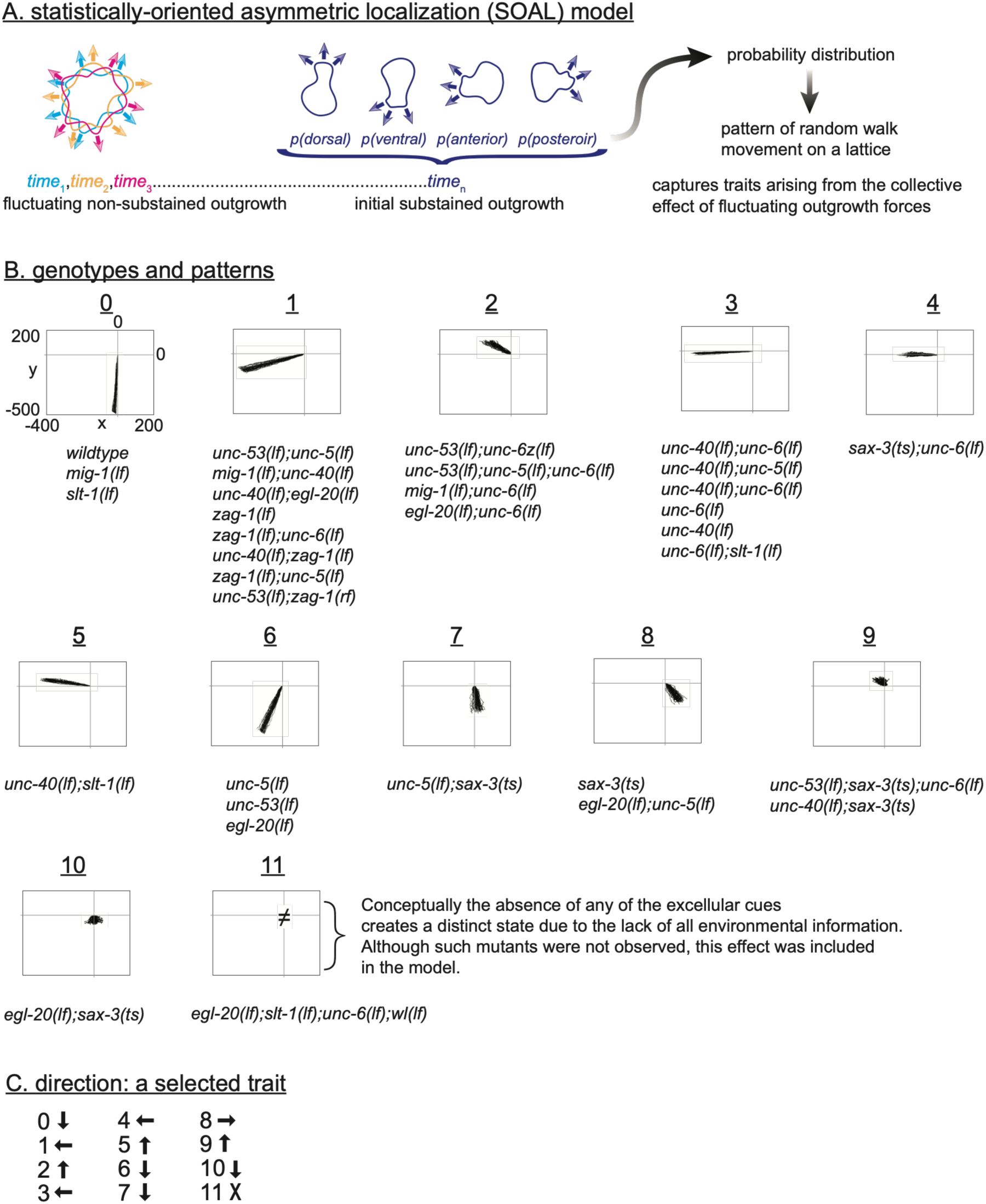
The statistically oriented asymmetric localization (SOAL) model and random walk simulations define discrete morphological macrostates. (A) The SOAL model conceptualizes axon outgrowth as a stochastically driven process. Left: In an equilibrium state lacking guidance cues, outgrowth probability is uniformly distributed in all directions. Middle: Extracellular guidance cues induce asymmetric, probability-based distributions of receptors and signaling molecules on the growth cone membrane, biasing outgrowth directionality. Right: These empirically derived probability distributions are utilized to generate biased random walk patterns on a lattice, providing a physical model for how fluctuations in cytoskeletal outgrowth forces translate into directed movements over time (visualized as 25 superimposed walks of 500 steps). (B) Phenotypic convergence of diverse genetic backgrounds into distinct topological patterns. Biased random walks were simulated using data from 33 different animal genotypes, including wild-type, single, and combinatorial mutants. Despite the underlying stochasticity of the force-producing machinery, the resulting spatial trajectories reliably merge into 12 discrete, stable topological patterns. These 12 morphological categories define the system’s observable macrostates. (C) Directional traits associated with each morphological macrostate. Based on a visual comparison of the distinct random walk topologies generated in (B), each macrostate is assigned a primary macroscopic outgrowth direction (e.g., anterior, ventral), linking the statistical properties of the genetic network to the emergent physical trajectory.

Despite the stochastic nature of the individual steps, the patterns revealed a striking phenotypic convergence. That is, the single and combinatorial mutants can be merged into 11 distinct topological patterns of outgrowth (Figure 2B). (For modeling, we additionally included an isotropic “null” macrostate representing unbiased outgrowth in the absence of any extracellular cues.) The observed convergence indicates that the genetic network modulates the statistical parameters that control the force-generating machinery of outgrowth movement. The genetic state constrains the randomness of movement and, in doing so, integrates the various signaling inputs into a finite set of stable topological outcomes. Further, this result indicates that local energy is expended to drive a statistically biased directed motion, which is consistent with the dynamics of physical systems operating far from equilibrium.

### Construction of a Computational Proxy for the Genetic Regulatory Network

While an unambiguous reconstruction of the molecular network linking gene function to force distribution would be ideal, the complexity of epistasis and redundancy makes this currently intractable. However, for modeling pathfinding, such a reconstruction might not be necessary. Instead, a predictive model of the system that could take input and accurately predict outgrowth behavior would be useful. One challenge to developing a model, however, is that linking the combinatorial genotypes to the specific force distributions requires non-linear logic. Artificial Neural Networks (ANNs or Multi-Layer Perceptrons) are very good at approximating complex, unknown, and high-dimensional relationships. An Artificial Neural Network (MLP, Multi-Layer Perceptron) was trained to serve as a functional proxy for the biological regulatory system. The MLP was designed to map an 11-dimensional binary input vector representing the loss-of-function (0) or wild-type (1) state of each gene to one of the distinct morphological patterns identified in the random walk analysis. To prevent the model from defaulting to the most common biological phenotypes, random oversampling was utilized, ensuring that rare yet topologically critical morphological attractors were weighted equally during training.

It is observed that no single MLP model could fully capture the complexity of genetic interactions. Over 1,000 independent models were initialized with different random weights and trained. When evaluated on an unseen test set, many models achieved high overall accuracy but failed to predict specific subsets of possible genotypes. However, an input that confused one model could be correctly classified by another. This observation implies that the regulatory logic is modular and distributed and that it is difficult to package into a single set of synaptic weights. To construct a comprehensive MLP model that no single model could capture in isolation, five models with complementary error profiles were selected and integrated into a single ensemble architecture. This ensemble processes inputs in parallel and averages their probability distributions.

To validate the predictive accuracy of the trained MLP, we used the random walk framework to assess its ability to replicate the macroscopic outgrowth biases observed in vivo. We calculated the expected two-dimensional displacement of a random walk based on the probabilities of outgrowth in the four cardinal directions (anterior, posterior, dorsal, and ventral). We compared displacements using probabilities from in vivo observations with those from the MLP’s output. For comparative scaling, the MLP’s raw output probabilities were normalized to the baseline probabilities of the experimental observations.

The trained MLP serves as a computational proxy for the cellular steering machinery. The model predicts the dominant macrostate when given the inputs for cellular gene functions and extracellular factors, and, because macrostates are associated with directional movement traits derived from random walk patterns (Figure 2C), the MLP output can be used to predict directional movements. We find that the random walk displacements generated by the computational proxy closely match the distribution of displacements derived directly from experimental data (Figure 3, Table). To test whether the MLP captured known genetic interactions of the GSN, we performed an *in silico* epistasis analysis. We evaluated all 2^3^ = 8 knockout states of unc-40, unc-5, and unc-6, and combined each state with an additional single knockout of one of eight genes (egl-20, mig-1, sax-3, slt-1, unc-53, wl, wr, or zag-1), yielding 77 input vectors. As a specific, measurable phenotype, we calculated the spatial displacement (Δ*d*) for each genetic perturbation, defined as the Euclidean distance between the 2D spatial coordinates predicted by the MLP for the baseline genetic background and those for the modified background containing the secondary knockout. This epistasis analysis focuses on the module comprising the UNC-6 (netrin) ligand and its receptors UNC-40 (DCC) and UNC-5 (UNC5). The genetic and molecular interactions among the components of this module have been well studied, and this module is critical for the wild-type straight-line movement (Macrostate 0). Because loss of *unc-6* or *unc-40* function results in straight-line movement in a different direction (Macrostate 3), we reason that the magnitude of spatial displacement reflects to some degree the volatility of the GNS. The magnitude of change in displacements across different gene combinations (epistatic volatility) is visualized in a heatmap (Figure 4). It is worth noting the distinction between the qualitative grouping of *in vivo* outgrowth in Figure 2B and the precise quantitative displacements generated by the MLP in Figure 4. The MLP reveals nuances that human observers miss by categorical binning.

**Figure 3.**
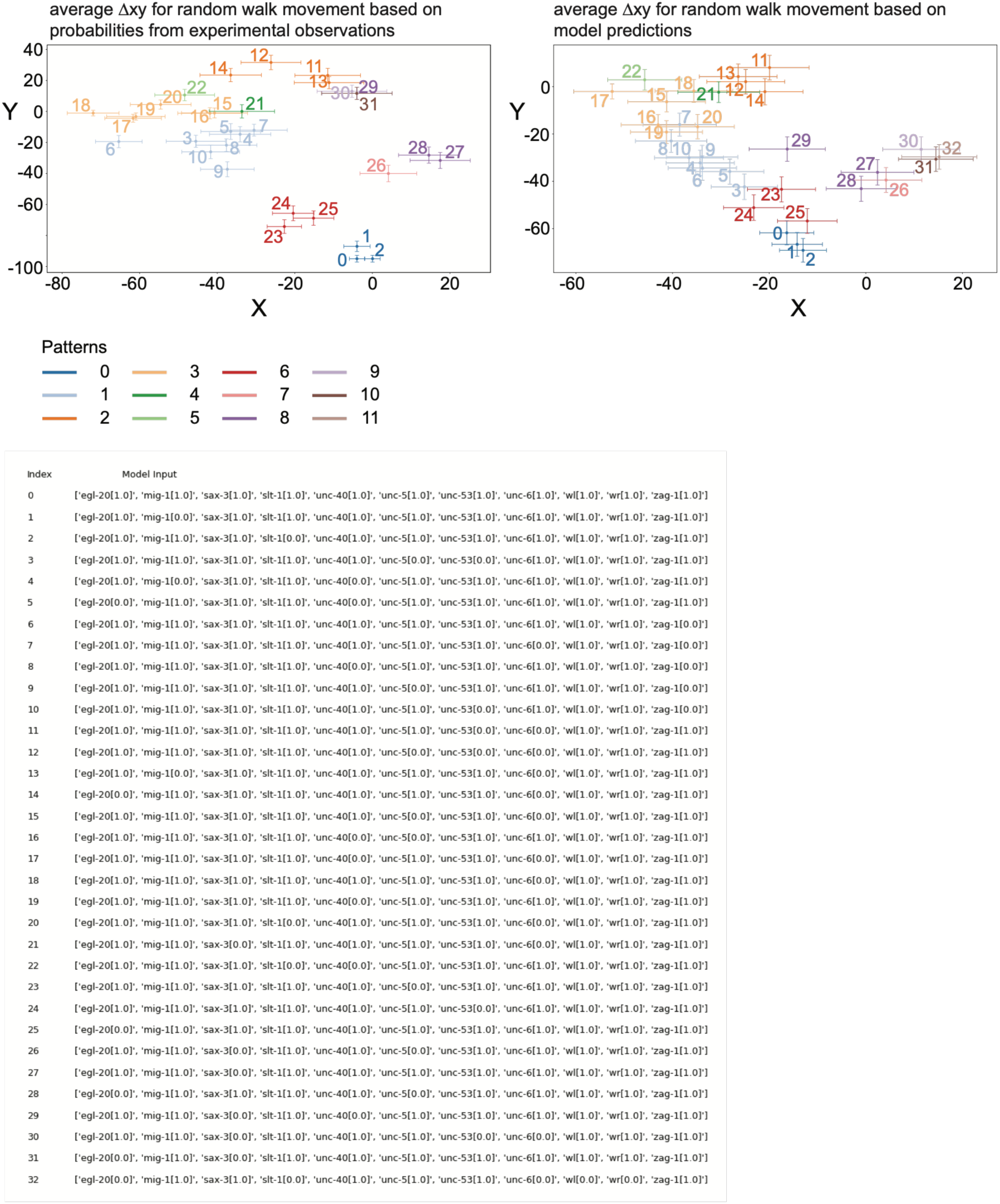
The trained Multi-Layer Perceptron (MLP) model accurately predicts empirical random walk displacements and macrostate topologies. (Top) Scatter plots comparing the average spatial displacements (Δx, Δy) of random walk simulations generated from 33 different genetic combinations. Displacements derived directly from experimental observations are compared with those predicted by the MLP computational proxy. The model’s predicted probability distributions were normalized to the empirical dataset to allow a direct comparison. The similar spatial clustering of specific gene combinations indicates that the MLP model accurately predicts the dominant directional macrostate derived from the complex genetic perturbations. The MLP successfully encodes the combinatorial logic of the Guidance Signaling Network (GSN). Data points are numbered by genotype and color-coded to indicate the specific morphological macrostate to which each belongs (defined in Figure 2B). (Bottom) Table detailing the specific 11-component genetic input vectors (wild-type vs. null states) corresponding to the numbered indices labeled in the scatter plots above.

**Figure 4.**
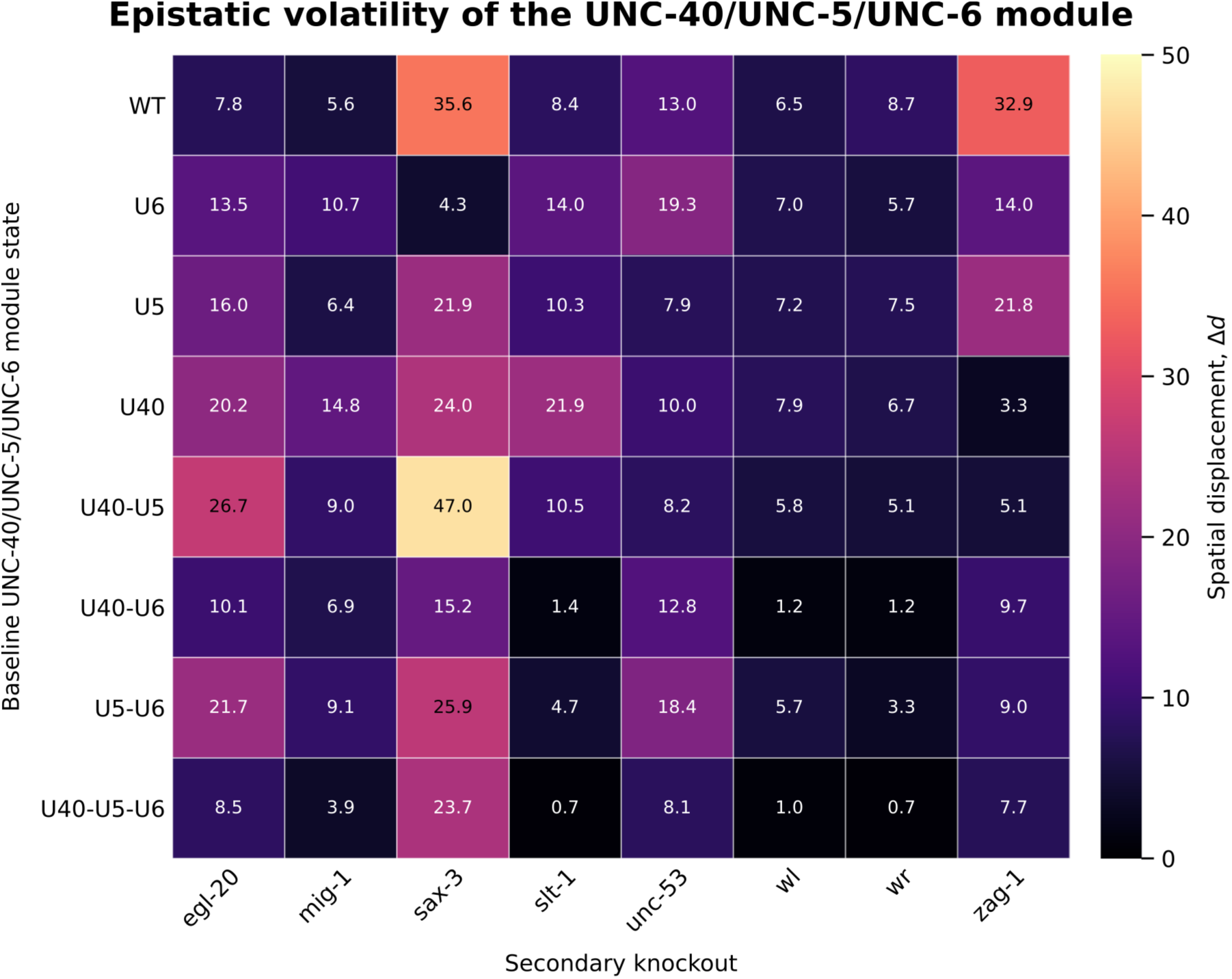
Epistatic Volatility of the U40-U5-U6 Module Across Genetic Backgrounds. Heatmap displaying the spatial displacement (Δ*d*) triggered by the introduction of secondary pathway knockouts (columns) into *unc-40* (U40)*, unc-5* (U5), and *unc-6* (U6) knockout combinations (rows). Δ*d* represents the Euclidean distance between the spatial coordinates predicted by the trained MLP for the baseline mutant and the modified genotype.

The *in silico* epistasis analysis tests whether the MLP captures the rules underlying the genetic relationships required to predict phenotypes when the model is given data it was not trained on. Previously, a stochastic framework for axon guidance was used to uncover genetic interactions among these genes. It was found that the interactions involve highly nonlinear epistatic suppression (24–26). A trained MLP model that failed to learn the underlying patterns and instead simply memorized the data (overfitting) would make erratic, irrational predictions on new data, interpreting random noise as epistatic interactions. This could be revealed by comparing the observed genetic interactions to those predicted by the model. In the epistasis test we performed, 48 of the 77 input vectors are data not used in training the model.

The epistatic volatility heatmap indicates that the MLP model learned the non-linear genetic rules of the GSN. For example, *zag-1* acts as a master transcriptional regulator (30, 31). Loss of *zag-1* in the wild-type background causes a major displacement (Δ*d* = 32.9), with loss of *unc-40,* the displacement is small (Δ*d* = 3.3). This rescue is not seen with the loss of *unc-5* (Δ*d* = 21.8).

Finally, the loss of *unc-6* in the loss of *zag-1* background reduces the severity (Δ*d* = 32.9 to Δ*d* = 14.0), and the loss of *unc-6, unc-40,* and *unc-5* is low (Δ*d* = 7.7). These results are consistent with evidence that UNC-40 drives stochastic fluctuations in outgrowth activity, whereas the UNC-6 ligand biases the orientation of UNC-40 asymmetry (32). Also, with evidence that UNC-5 establishes statistically oriented, restrictive spatial boundaries on outgrowth (25). Our interpretation is that the model correctly identifies that, without ZAG-1 activity, stochastic fluctuations in outgrowth driven by UNC-40 are not properly regulated, thereby diminishing UNC-40’s effectiveness in driving outgrowth in any direction. This chaotic outgrowth is removed with the loss of UNC-40. UNC-5 does not have this same activity; together with UNC-6, it restricts the spatial boundaries of the stochastic UNC-40-mediated outgrowth. Similarly, the results in the loss of *unc-53* background and in the loss of *slit-1* backgrounds also highlight the role of UNC-40 as a biological engine that intrinsically generates stochastic fluctuations in outgrowth and that UNC-6 and UNC-5 modify this activity. In the loss of *unc-53* background, the loss of *unc-6,* or the loss of *unc-5* and *unc-6,* resulted in functionally similar epistatic volatility (Δ*d* = 19.3 vs. 18.4), reflecting UNC-5’s dependence on its ligand. In contrast, the loss of *unc-40* in the same *unc-6* null background significantly altered the network state (Δ*d* = 12.8). The volatility induced by the loss of *slt-1* in the *unc-6* null background (Δ*d* = 14.0) was nearly completely rescued by the removal of UNC-40 (Δ*d* = 1.40). The model also captures the interwoven activities of UNC-40 and SAX-3 that produce directional movement. Previous *in vivo* studies examining axon guidance as a stochastic process showed that both act in establishing a directional bias. The evidence suggests that SAX-3 is required to induce the UNC-40 asymmetric localization and that this induction is a necessity for the process that determines the probability of where UNC-40 localizes in response to an extracellular cue (24). The MLP results support this interpretation. It predicts a large spatial deviation when sax-3 was removed in a wild-type background (Δ*d* = 35.6). However, in an unc-6 null background, loss of unc-6 and sax-3 has little effect (Δ*d* = 4.3).

Finally, the disparity between categorical macrostate groupings and quantitative epistatic volatility indicates the MLP avoided overfitting. During training, both the unc-40; zag-1 and unc-6; zag-1 mutants were categorized into the same phenotypic basin of attraction (Macrostate 1). An overfit model, acting as a discrete lookup table, would map both inputs to identical spatial coordinates, resulting in equivalent Δ*d* displacements. Instead, the MLP predicted highly divergent epistatic volatilities for zag-1 loss in the unc-40 (Δ*d* = 3.3) versus unc-6 (Δ*d* = 14.0) backgrounds. Taken together, this *in silico* epistasis analysis indicates that the MLP did not overfit the data; it generalized to learn underlying, fundamental patterns and relationships among the genes.

### Simulation of Axon Trajectories via Maximum Entropy Steering

We adapted the MLP to map the local stability landscape of the growth cone at every spatial coordinate along a trajectory’s pathway. At each position, the MLP evaluates the 11 single-scalar values of the GSN, which represent cellular gene functions and extracellular factors. Standard artificial neural networks typically produce a single, fixed prediction for a given input. However, biological networks are inherently noisy and subject to stochastic fluctuations. In fact, the inspiration for exploring the GSN as a stochastic system came from evidence that the direction and timing of UNC-6 (netrin) mediated guidance is dependent on stochastic fluctuations in intracellular UNC-40 (DCC) outgrowth activity (26). We imagine that multiple cues act concurrently on spatially distinct membrane subdomains and drive rapid reconfiguration of receptor occupancy and downstream effectors. To emulate this instant-to-instant variability, we used the base state, together with a combinatorial dropout analysis that systematically evaluates a biologically constrained ensemble of null-mutation variants. The model evaluates all single-, double-, and triple-loss-of-function combinations by setting the corresponding gene inputs to zero. The perturbation ensemble was restricted to a set of 232 total conditions (231 perturbations; or 232 total conditions including the baseline) because genetic experiments indicate that loss-of-function mutations in four or more of these genes are typically lethal, or because we considered them to lack developmental significance (such as when no values for extracellular cues were included).

To quantify the network’s thermodynamic properties during a simulated axon guidance, we calculated entropy at each spatial coordinate. Let *m* represent a specific combinatorial microstate of the Guidance Signaling Network (GSN) and *k* represent a distinct morphological macrostate (Figure 2B). Because biological networks exhibit profound degeneracy, multiple microstates can belong to the same macrostate. Using the trained Multi-Layer Perceptron (MLP) as a functional proxy, the raw probability *P*(*y* = *k*|*m*) that a given microstate produces a specific macrostate was determined. To assess the total distribution of available microstates for any given macrostate, these probabilities were normalized across the entire input ensemble to yield

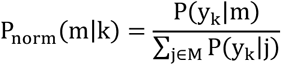

Finally, the Shannon entropy (*H_k_*) of each of the 12 morphological macrostate was calculated using the natural logarithm. To prevent mathematically undefined values, any microstates where *P_norm_* = 0 were filtered out and excluded from the summation,

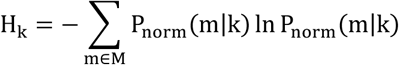

In this biophysical framework, *H_k_* explicitly quantifies the network degeneracy underlying a physical behavior.

- Low Macrostate Entropy: a macrostate with a low *H_k_* value represents a deep, energy-minimized attractor; the growth cone is restricted to a highly specific set of microstates. It is resistant to minor genetic or environmental fluctuations.
- High Macrostate Entropy: a macrostate with a high *H_k_* value reflects an expansive attractor landscape. The growth cone can readily swap between many underlying microstates to maintain its trajectory. A high *H_k_* denotes high plasticity, where rapid switching among multiple morphological macrostates is supported.

The simulation utilizes these 12 discrete entropy values as a steering algorithm. At each step, the algorithm (representing a decision-making framework of a growth cone) samples the neighboring coordinates and computes the macrostate entropy for all possible directions.

Movement is based on the directional behavior trait associated with the macrostate exhibiting the maximum entropy. When the MLP model is initialized with baseline wild-type parameters derived from experimental observations (all set to 1.0, the wild-type values), the simulation accurately reproduces the wild-type trajectory of the HSN neuron, which in vivo migrates ventrally to the ventral midline (Figure 5A). Historically, this migration has been considered growth cone chemotaxis, that is, the growth cone ‘sensing’ a gradient and directing movement along it. For the HSN growth cone, movement is towards the source of UNC-6 (netrin), which is secreted by ventral midline cells. Strikingly, the simulation reproduces the wild-type HSN trajectory without involving a gradient-following behavior. Instead, movement emerges from selecting directions that maximize local signaling entropy. This result suggests that growth cone guidance could involve an active search for states of high signaling sensitivity, rather than relying solely on deterministic chemotaxis.

**Figure 5.**
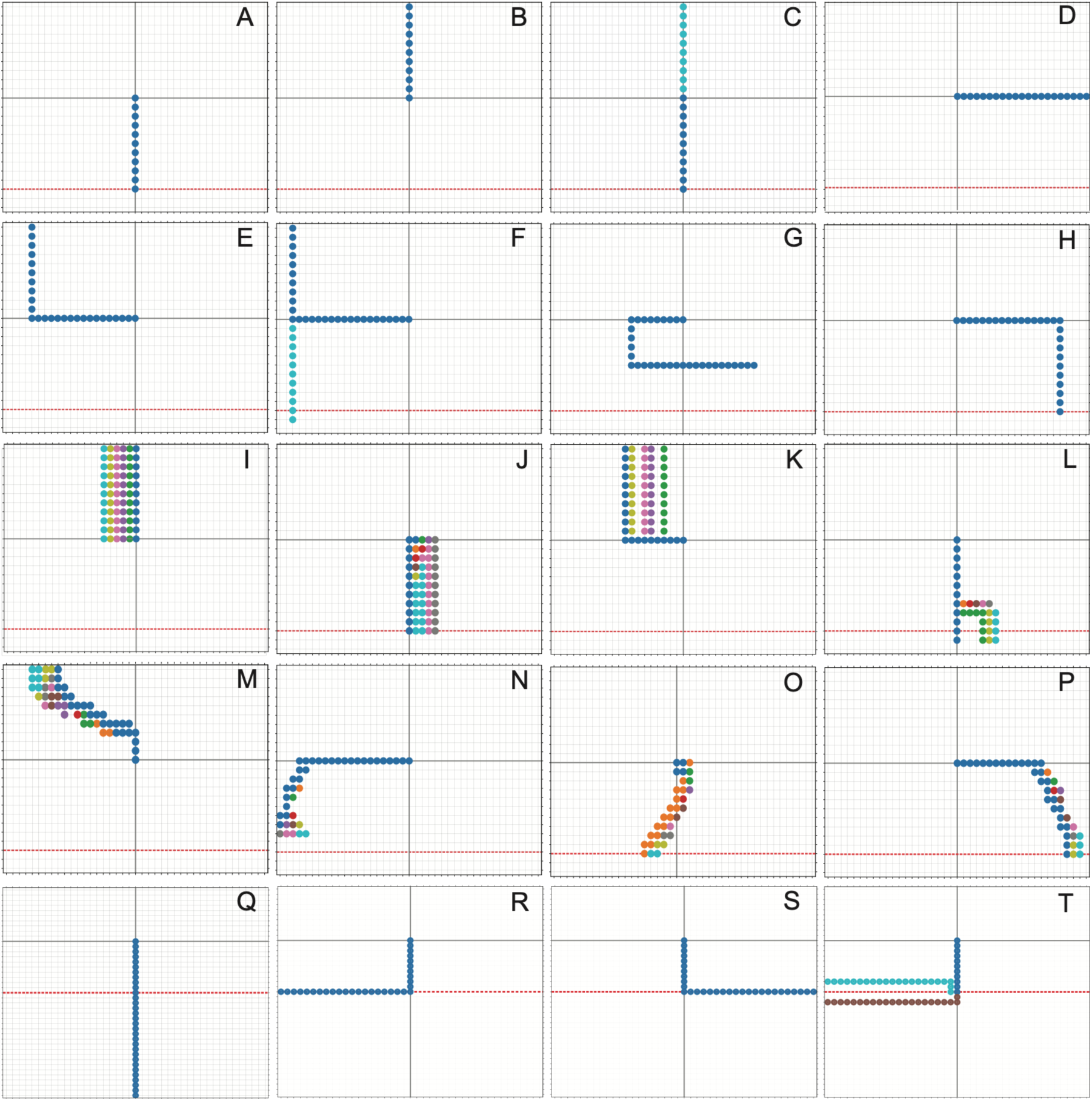
Complex neural trajectories and choice point navigation emerge from tuning intrinsic gene states within a static environment. (A-P) Monte Carlo simulations sampling the parameter space of intrinsic gene functions yield highly diverse pathfinding topologies within a single, constant extracellular environment. Panel A displays the simulated wild-type HSN trajectory driven by baseline intrinsic values (all equal to 1.0), resulting in a ventral trajectory toward the midline and modeling HSN growth cone migration in vivo. Panels B-P demonstrate diverse divergent phenotypes resulting from altered intrinsic states. Simulations are mapped on the right side of the anatomical axis, i.e., anterior is to the left. (Q-T) Simulations of navigational decisions at the ventral midline choice point. Trajectories initiate with wild-type intrinsic parameters, driving the axon to the midline. Upon reaching the midline, new values for the intrinsic gene states are applied to simulate contact-induced alterations to the internal signaling network. Tuning these internal states enables the growth cone to execute complex choice point behaviors, including contralateral crossing (Q), longitudinal anterior turning (R), longitudinal posterior turning (S), and bilateral branching (T). Grid Orientation & Markers: In all panels, anterior is oriented to the left. The intersection of the solid gray lines indicates the starting coordinate where all local extracellular cue values (UNC-6, SLT-1, EGL-20, WL) are equal to 1.0. The red dashed line denotes the ventral midline, corresponding to the spatial coordinates of maximum UNC-6 and minimum SLT-1 values. Values used for the plots are given in Table 1.

**Table 1.**
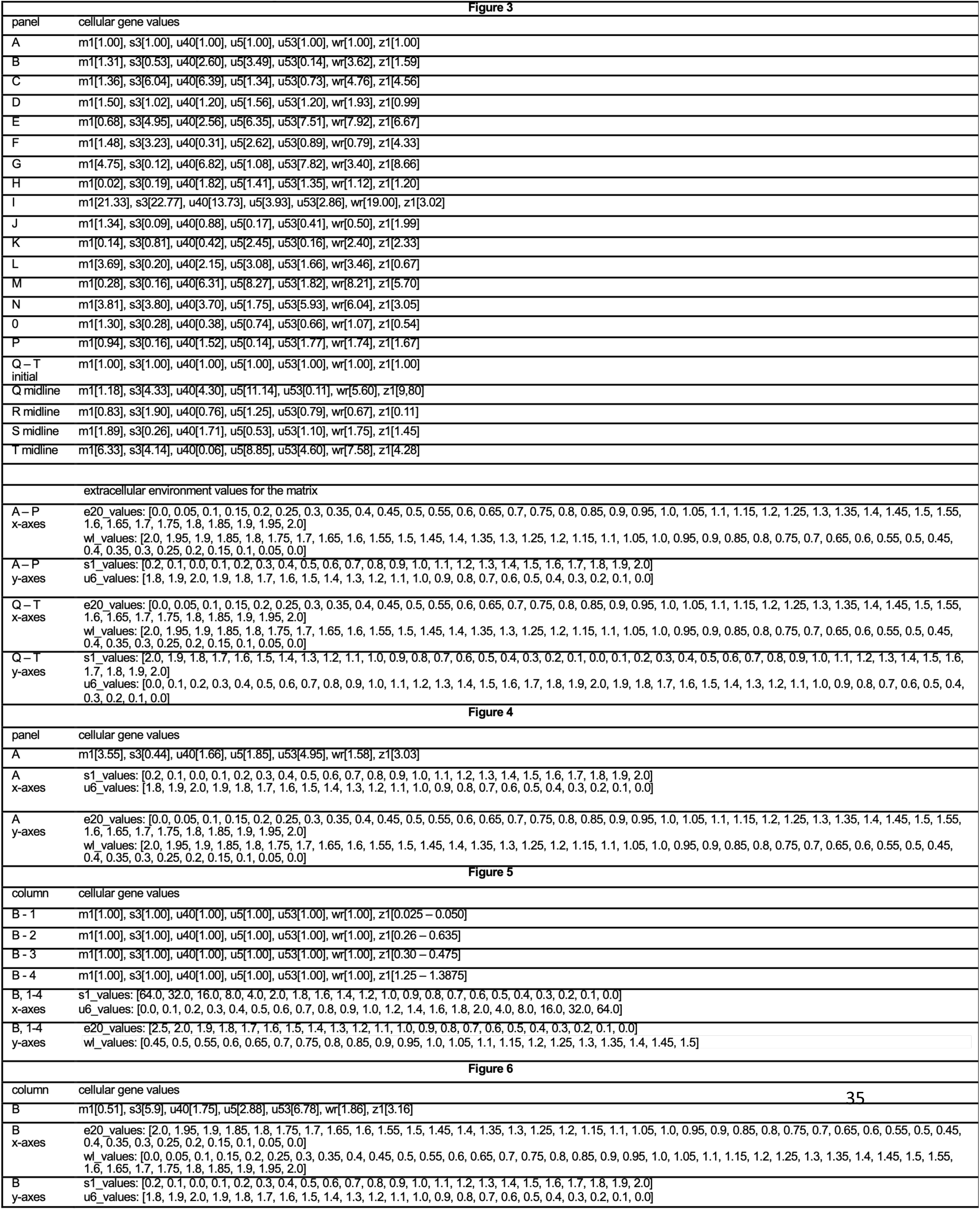
Values used for plots in Figures 3,4,5, and 6.

| panel | cellular gene values |
| --- | --- |
| A | m1[1.00], s3[1.00], u40[1.00], u5[1.00], u53[1.00], wr[1.00], z1[1.00] |
| B | m1[1.31], s3[0.53], u40[2.60], u5[3.49], u53[0.14], wr[3.62], z1[1.59] |
| C | m1[1.36], s3[6.04], u40[6.39], u5[1.34], u53[0.73], wr[4.76], z1[4.56] |
| D | m1[1.50], s3[1.02], u40[1.20], u5[1.56], u53[1.20], wr[1.93], z1[0.99] |
| E | m1[0.68], s3[4.95], u40[2.56], u5[6.35], u53[7.51], wr[7.92], z1[6.67] |
| F | m1[1.48], s3[3.23], u40[0.31], u5[2.62], u53[0.89], wr[0.79], z1[4.33] |
| G | m1[4.75], s3[0.12], u40[6.82], u5[1.08], u53[7.82], wr[3.40], z1[8.66] |
| H | m1[0.02], s3[0.19], u40[1.82], u5[1.41], u53[1.35], wr[1.12], z1[1.20] |
| I | m1[21.33], s3[22.77], u40[13.73], u5[3.93], u53[2.86], wr[19.00], z1[3.02] |
| J | m1[1.34], s3[0.09], u40[0.88], u5[0.17], u53[0.41], wr[0.50], z1[1.99] |
| K | m1[0.14], s3[0.81], u40[0.42], u5[2.45], u53[0.16], wr[2.40], z1[2.33] |
| L | m1[3.69], s3[0.20], u40[2.15], u5[3.08], u53[1.66], wr[3.46], z1[0.67] |
| M | m1[0.28], s3[0.16], u40[6.31], u5[8.27], u53[1.82], wr[8.21], z1[5.70] |
| N | m1[3.81], s3[3.80], u40[3.70], u5[1.75], u53[5.93], wr[6.04], z1[3.05] |
| O | m1[1.30], s3[0.28], u40[0.38], u5[0.74], u53[0.66], wr[1.07], z1[0.54] |
| P | m1[0.94], s3[0.16], u40[1.52], u5[0.14], u53[1.77], wr[1.74], z1[1.67] |
| Q – T<br>initial | m1[1.00], s3[1.00], u40[1.00], u5[1.00], u53[1.00], wr[1.00], z1[1.00] |
| Q midline | m1[1.18], s3[4.33], u40[4.30], u5[11.14], u53[0.11], wr[5.60], z1[9.80] |
| R midline | m1[0.83], s3[1.90], u40[0.76], u5[1.25], u53[0.79], wr[0.67], z1[0.11] |
| S midline | m1[1.89], s3[0.26], u40[1.71], u5[0.53], u53[1.10], wr[1.75], z1[1.45] |
| T midline | m1[6.33], s3[4.14], u40[0.06], u5[8.85], u53[4.60], wr[7.58], z1[4.28] |
| extracellular environment values for the matrix |  |
| A – P<br>x-axes | e20_values: [0.0, 0.05, 0.1, 0.15, 0.2, 0.25, 0.3, 0.35, 0.4, 0.45, 0.5, 0.55, 0.6, 0.65, 0.7, 0.75, 0.8, 0.85, 0.9, 0.95, 1.0, 1.05, 1.1, 1.15, 1.2, 1.25, 1.3, 1.35, 1.4, 1.45, 1.5, 1.55, 1.6, 1.65, 1.7, 1.75, 1.8, 1.85, 1.9, 1.95, 2.0]<br>wl_values: [2.0, 1.95, 1.9, 1.85, 1.8, 1.75, 1.7, 1.65, 1.6, 1.55, 1.5, 1.45, 1.4, 1.35, 1.3, 1.25, 1.2, 1.15, 1.1, 1.05, 1.0, 0.95, 0.9, 0.85, 0.8, 0.75, 0.7, 0.65, 0.6, 0.55, 0.5, 0.45, 0.4, 0.35, 0.3, 0.25, 0.2, 0.15, 0.1, 0.05, 0.0] |
| A – P<br>y-axes | s1_values: [0.2, 0.1, 0.0, 0.1, 0.2, 0.3, 0.4, 0.5, 0.6, 0.7, 0.8, 0.9, 1.0, 1.1, 1.2, 1.3, 1.4, 1.5, 1.6, 1.7, 1.8, 1.9, 2.0]<br>u6_values: [1.8, 1.9, 2.0, 1.9, 1.8, 1.7, 1.6, 1.5, 1.4, 1.3, 1.2, 1.1, 1.0, 0.9, 0.8, 0.7, 0.6, 0.5, 0.4, 0.3, 0.2, 0.1, 0.0] |
| Q – T<br>x-axes | e20_values: [0.0, 0.05, 0.1, 0.15, 0.2, 0.25, 0.3, 0.35, 0.4, 0.45, 0.5, 0.55, 0.6, 0.65, 0.7, 0.75, 0.8, 0.85, 0.9, 0.95, 1.0, 1.05, 1.1, 1.15, 1.2, 1.25, 1.3, 1.35, 1.4, 1.45, 1.5, 1.55, 1.6, 1.65, 1.7, 1.75, 1.8, 1.85, 1.9, 1.95, 2.0]<br>wl_values: [2.0, 1.95, 1.9, 1.85, 1.8, 1.75, 1.7, 1.65, 1.6, 1.55, 1.5, 1.45, 1.4, 1.35, 1.3, 1.25, 1.2, 1.15, 1.1, 1.05, 1.0, 0.95, 0.9, 0.85, 0.8, 0.75, 0.7, 0.65, 0.6, 0.55, 0.5, 0.45, 0.4, 0.35, 0.3, 0.25, 0.2, 0.15, 0.1, 0.05, 0.0] |
| Q – T<br>y-axes | s1_values: [2.0, 1.9, 1.8, 1.7, 1.6, 1.5, 1.4, 1.3, 1.2, 1.1, 1.0, 0.9, 0.8, 0.7, 0.6, 0.5, 0.4, 0.3, 0.2, 0.1, 0.0, 0.1, 0.2, 0.3, 0.4, 0.5, 0.6, 0.7, 0.8, 0.9, 1.0, 1.1, 1.2, 1.3, 1.4, 1.5, 1.6, 1.7, 1.8, 1.9, 2.0]<br>u6_values: [0.0, 0.1, 0.2, 0.3, 0.4, 0.5, 0.6, 0.7, 0.8, 0.9, 1.0, 1.1, 1.2, 1.3, 1.4, 1.5, 1.6, 1.7, 1.8, 1.9, 2.0, 1.9, 1.8, 1.7, 1.6, 1.5, 1.4, 1.3, 1.2, 1.1, 1.0, 0.9, 0.8, 0.7, 0.6, 0.5, 0.4, 0.3, 0.2, 0.1, 0.0] |

**Figure 4**
| panel | cellular gene values |
| --- | --- |
| A | m1[3.55], s3[0.44], u40[1.66], u5[1.85], u53[4.95], wr[1.58], z1[3.03] |
| A<br>x-axes | s1_values: [0.2, 0.1, 0.0, 0.1, 0.2, 0.3, 0.4, 0.5, 0.6, 0.7, 0.8, 0.9, 1.0, 1.1, 1.2, 1.3, 1.4, 1.5, 1.6, 1.7, 1.8, 1.9, 2.0]<br>u6_values: [1.8, 1.9, 2.0, 1.9, 1.8, 1.7, 1.6, 1.5, 1.4, 1.3, 1.2, 1.1, 1.0, 0.9, 0.8, 0.7, 0.6, 0.5, 0.4, 0.3, 0.2, 0.1, 0.0] |
| A<br>y-axes | e20_values: [0.0, 0.05, 0.1, 0.15, 0.2, 0.25, 0.3, 0.35, 0.4, 0.45, 0.5, 0.55, 0.6, 0.65, 0.7, 0.75, 0.8, 0.85, 0.9, 0.95, 1.0, 1.05, 1.1, 1.15, 1.2, 1.25, 1.3, 1.35, 1.4, 1.45, 1.5, 1.55, 1.6, 1.65, 1.7, 1.75, 1.8, 1.85, 1.9, 1.95, 2.0]<br>wl_values: [2.0, 1.95, 1.9, 1.85, 1.8, 1.75, 1.7, 1.65, 1.6, 1.55, 1.5, 1.45, 1.4, 1.35, 1.3, 1.25, 1.2, 1.15, 1.1, 1.05, 1.0, 0.95, 0.9, 0.85, 0.8, 0.75, 0.7, 0.65, 0.6, 0.55, 0.5, 0.45, 0.4, 0.35, 0.3, 0.25, 0.2, 0.15, 0.1, 0.05, 0.0] |

**Figure 5**
| column | cellular gene values |
| --- | --- |
| B - 1 | m1[1.00], s3[1.00], u40[1.00], u5[1.00], u53[1.00], wr[1.00], z1[0.025 – 0.050] |
| B - 2 | m1[1.00], s3[1.00], u40[1.00], u5[1.00], u53[1.00], wr[1.00], z1[0.26 – 0.635] |
| B - 3 | m1[1.00], s3[1.00], u40[1.00], u5[1.00], u53[1.00], wr[1.00], z1[0.30 – 0.475] |
| B - 4 | m1[1.00], s3[1.00], u40[1.00], u5[1.00], u53[1.00], wr[1.00], z1[1.25 – 1.3875] |
| B, 1-4<br>x-axes | s1_values: [64.0, 32.0, 16.0, 8.0, 4.0, 2.0, 1.8, 1.6, 1.4, 1.2, 1.0, 0.9, 0.8, 0.7, 0.6, 0.5, 0.4, 0.3, 0.2, 0.1, 0.0]<br>u6_values: [0.0, 0.1, 0.2, 0.3, 0.4, 0.5, 0.6, 0.7, 0.8, 0.9, 1.0, 1.2, 1.4, 1.6, 1.8, 2.0, 4.0, 8.0, 16.0, 32.0, 64.0] |
| B, 1-4<br>y-axes | e20_values: [2.5, 2.0, 1.9, 1.8, 1.7, 1.6, 1.5, 1.4, 1.3, 1.2, 1.1, 1.0, 0.9, 0.8, 0.7, 0.6, 0.5, 0.4, 0.3, 0.2, 0.1, 0.0]<br>wl_values: [0.45, 0.5, 0.55, 0.6, 0.65, 0.7, 0.75, 0.8, 0.85, 0.9, 0.95, 1.0, 1.05, 1.1, 1.15, 1.2, 1.25, 1.3, 1.35, 1.4, 1.45, 1.5] |

**Figure 6**
| column | cellular gene values |
| --- | --- |
| B | m1[0.51], s3[5.9], u40[1.75], u5[2.88], u53[6.78], wr[1.86], z1[3.16] |
| B<br>x-axes | e20_values: [2.0, 1.95, 1.9, 1.85, 1.8, 1.75, 1.7, 1.65, 1.6, 1.55, 1.5, 1.45, 1.4, 1.35, 1.3, 1.25, 1.2, 1.15, 1.1, 1.05, 1.0, 0.95, 0.9, 0.85, 0.8, 0.75, 0.7, 0.65, 0.6, 0.55, 0.5, 0.45, 0.4, 0.35, 0.3, 0.25, 0.2, 0.15, 0.1, 0.05, 0.0]<br>wl_values: [0.0, 0.05, 0.1, 0.15, 0.2, 0.25, 0.3, 0.35, 0.4, 0.45, 0.5, 0.55, 0.6, 0.65, 0.7, 0.75, 0.8, 0.85, 0.9, 0.95, 1.0, 1.05, 1.1, 1.15, 1.2, 1.25, 1.3, 1.35, 1.4, 1.45, 1.5, 1.55, 1.6, 1.65, 1.7, 1.75, 1.8, 1.85, 1.9, 1.95, 2.0] |
| B<br>y-axes | s1_values: [0.2, 0.1, 0.0, 0.1, 0.2, 0.3, 0.4, 0.5, 0.6, 0.7, 0.8, 0.9, 1.0, 1.1, 1.2, 1.3, 1.4, 1.5, 1.6, 1.7, 1.8, 1.9, 2.0]<br>u6_values: [1.8, 1.9, 2.0, 1.9, 1.8, 1.7, 1.6, 1.5, 1.4, 1.3, 1.2, 1.1, 1.0, 0.9, 0.8, 0.7, 0.6, 0.5, 0.4, 0.3, 0.2, 0.1, 0.0] |

### Emergence of Complex Neural Architectures via Tuning of Intrinsic Gene States

Having established that entropic phase transitions can steer growth cones, we asked whether this mechanism could account for the morphological diversity of outgrowth patterns observed in vivo. The growth cones of distinct neuronal classes navigate the same global extracellular environment but adopt radically different trajectories. We hypothesized that this diversity arises because differences in the state of growth cone intrinsic cellular gene functions uniquely warp each cell type’s entropic landscape.

To test this, the spatial matrix that simulates extracellular gradients (UNC-6, SLT-1, EGL-20, WL) was held constant, and a Monte Carlo simulation was used to stochastically sample the parameter space of the intrinsic cellular gene functions. The algorithm successfully identified distinct intrinsic states that generated a vast array of trajectory patterns (Figure 5, Panels A-P). By merely altering the values for growth cone cellular gene inputs, the same static environment produced diverse behaviors, including turning, branching, orthogonal, and periodic movements.

We also applied this framework to model navigation at “choice points”. Choice points are intermediate locations where growth cones must make critical navigational decisions, such as turning or branching, to reach the final target. Early observation of axon guidance in the limb buds of embryonic grasshoppers indicated that guidepost cells were required for correct axon trajectories (33). The most intensively studied choice point is the ventral midline, where patterning is critical for bilateral organisms to coordinate the left and right halves of their bodies. At the ventral midline, growth cones frequently pause, interact with localized cues, and alter their trajectory (34–36). Contact with midline cues can trigger transcriptional and translational changes in the growth cone and shift its intrinsic signaling state, allowing new behaviors, such as crossing or turning.

We simulated this dynamic at the choice point using a two-phase approach. First, the growth cone is initialized with wild-type parameters, which drives it to the ventral midline (Figure 5, Panels Q-T). Upon reaching the midline, where UNC-6 and SLT-1 have their maximum and minimum values, respectively, a Monte Carlo simulation is applied to generate new intrinsic gene states. This mimics midline-induced signaling alterations. This localized tuning of the intrinsic state successfully enabled the growth cone to escape the midline attractor and execute complex, divergent navigational decisions. Depending on what intrinsic state was induced at the choice point, trajectories crossed the midline to the contralateral side (Figure 5Q), executed sharp anterior (Figure 5R) or posterior (Figure 5S) turns, or bifurcated and branched bilaterally (Figure 5T).

These results demonstrate that complex neural architectures do not strictly require highly complex or dynamically shifting extracellular gradients. Instead, modifications of intrinsic cellular gene states, which reposition the growth cone within the entropic landscape, are sufficient to generate a full spectrum of in vivo axon guidance behaviors, including directional plasticity, branching and bifurcations, bipolar outgrowths, periodic patterning, regional specification, and choice-point crossing.

### Directional Decisions Coincide with Entropic Phase Transitions

To better understand how the entropic landscape relates to turning behavior during outgrowth, we mapped stepwise the changes in macrostate entropy onto the corresponding turning events. A simulated trajectory was used that exhibits a complex navigational sequence. An initial anterior migration is followed by a ventral turn, a subsequent posterior turn, and a final ventral turn toward the midline (Figure 6A, labeled 1, 2, and 3, respectively). At each spatial coordinate (time step), the algorithm computes the Shannon entropy for all 12 morphological macrostates, yielding a profile of the competing macrostate influences within the system (Figure 6B).

**Figure 6.**
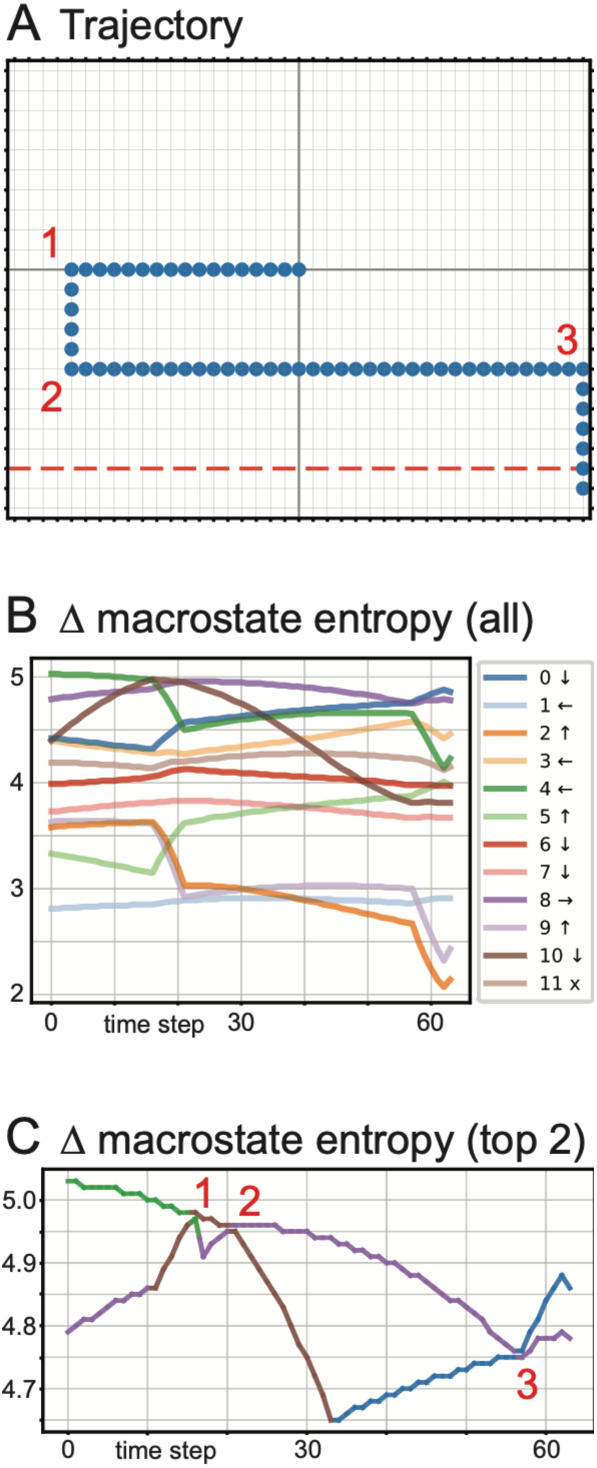
Entropic phase transitions drive axonal turning events. (A) A simulated 2D trajectory exhibiting complex pathfinding, including sequential turns mapped at points 1, 2, and 3. (B) Continuous tracking of the Macrostate Entropy (H_k_) for all 12 morphological macrostates across the duration of the trajectory (time steps). The legend indicates the color-coding for each macrostate and its corresponding directional drive (arrows). (C) Isolation of the top two highest entropy values at each time step. Color changes in the topmost line represent an entropic crossover, a phase transition where a new macrostate becomes the most volatile and assumes dominance. The time steps corresponding to these entropic crossovers directly align with the physical turning points (1, 2, and 3) in the spatial trajectory shown in Panel A. Grid Orientation & Markers: In all panels, anterior is oriented to the left. The intersection of the solid gray lines indicates the starting coordinate where all local extracellular cue values (UNC-6, SLT-1, EGL-20, WL) are equal to 1.0. The red dashed line denotes the ventral midline, corresponding to the spatial coordinates of maximum UNC-6 and minimum SLT-1 values. Values used for the plots are given in Table 1.

We plotted the entropies of the two most dominant macrostates at each step (Figure 6C). This visualization illustrates that the system is frequently engaged in a competition between two highly unstable, equiprobable states. During periods of straight, continuous migration (e.g., the initial anterior growth), the entropy of the dominant macrostate remains safely above its competitors.

However, as the growth cone moves through extracellular gradients and the resulting intracellular signaling changes, the stability landscape is gradually reshaped. A physical change in direction occurs only when the entropy of a competing macrostate surpasses that of the current dominant one. These entropic crossovers (Figure 6C, points 1, 2, and 3) occur precisely at the spatial turning points in the trajectory (Figure 6A, points 1, 2, and 3). These shifts correspond to entropic phase transitions in which the system reorganizes around the macrostate with the fastest-rising instability, thereby prompting a new orientation of the cytoskeletal network.

### Temporal Dynamics of a Transcription Factor Regulate the Sequence of Developmental Phase Transitions

Neuronal development requires temporal precision. The correct positioning of the cell body and the subsequent routing of its axon occur in sync with other developmental events in the animal. In the HSN neuron, the transcription factor ZAG-1 is temporally regulated and is required for both the anterior migration of the cell body and the subsequent ventral guidance of the axon (30, 31). To investigate how temporal regulation of intrinsic gene states controls the sequence of navigational events, we simulated trajectories when the cellular zag-1 input value (z1) was incrementally increased at each time step. This represents a progressive accumulation of the transcription factor during development.

We held the extracellular matrix constant and tested four distinct temporal expression profiles (Figure 7A, Rows 1-4). This approach yielded different outgrowth trajectories (Figure 7B) driven by the shifting entropic landscape of the top competing macrostates (Figure 7C).

- Sub-threshold Accumulation (Row 1): With z1 starting low and rising only modestly (from 0.025 to 0.05), the internal dynamics never crossed the threshold needed to exit the initial attractor basin. As a result, Macrostate 1 (“Anterior”) consistently exhibited the dominant macrostate entropy, and the simulated axon advanced anteriorly without any directional change.
- Premature Phase Transition (Row 4): When z1 expression was initialized at a high baseline (1.25-1.38), the system began in a post-transition state. Macrostate 0 (“Ventral”) overtook Macrostate 1 almost immediately at time step 1, causing the trajectory to abort anterior migration and drive prematurely toward the ventral midline.
- Sequenced Navigation (Row 3): With an intermediate profile (0.30-0.475), the system initially favored Macrostate 1, driving early anterior migration. However, as z1 accumulated, the entropic landscape shifted, triggering a clean phase transition at time step 4. Macrostate 0 became dominant, forcing a definitive 90-degree ventral turn, effectively separating the trajectory into two distinct developmental phases.
- Entropic Equilibrium and Branching (Row 2): A steep accumulation profile (0.26 to 0.635) pushed the system into a state of profound instability. At time step 4, the entropies of Macrostate 1 and Macrostate 0 reached perfect equilibrium. In this state, the growth cone has maximum plasticity. There is a physical bifurcation where one branch continued anteriorly, driven by Macrostate 1, while a secondary trajectory successfully escaped the initial attractor and traveled ventrally under the influence of Macrostate 0.

**Figure 7.**
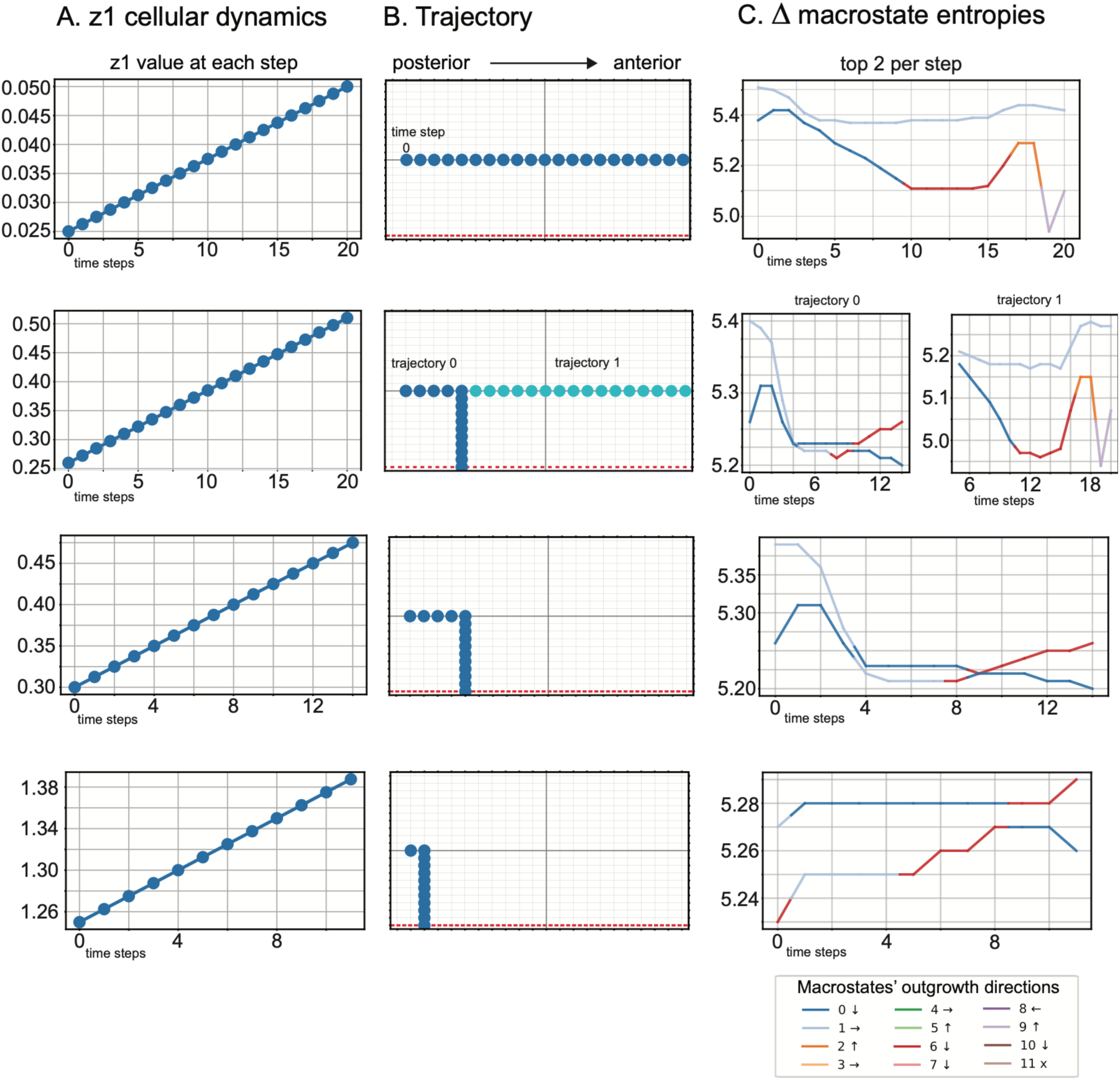
Temporal dynamics of the zag-1 transcription factor dictate the sequencing of entropic phase transitions. (A) Line plots illustrating the stepwise temporal accumulation of the cellular z1 (zag-1) parameter across four distinct simulation scenarios (Rows 1-4). (B) The time-dependent changes in the parameter generate the resulting 2D spatial trajectories. Row 1 yields continuous anterior migration; Row 2 results in a bifurcation at time step 4; Row 3 produces a sequenced anterior-to-ventral turn at time step 4; Row 4 exhibits a premature ventral turn at time step 1. (C) Tracking of the top two highest macrostate entropies at each time step. Crossover events between Macrostate 1 (cyan, anterior drive) and Macrostate 0 (orange, ventral drive) align precisely with the spatial turning points. In Row 2, the entropies of the two macrostates reach equilibrium at the turning point (time step 4), resolving into two distinct tracked trajectories (Trajectory 0 and Trajectory 1). Grid Orientation & Markers: In all rows of column B, the anterior direction is oriented to the right to align visually with the progression of the time axis. The intersection of the solid gray lines indicates the starting coordinate where all local extracellular cue values (UNC-6, SLT-1, EGL-20, WL) are equal to 1.0. The initial trajectory’s starting point is set relative to this intersection. The red dashed line denotes the ventral midline, corresponding to the spatial coordinates of maximum UNC-6 and minimum SLT-1 values. Values used for the plots are provided in Table 1.

These results indicate that temporal modulation of a transcription factor can drive the growth cone through a sequence of entropic phase transitions. The regulated expression of the transcription factor, in effect, acts as an intrinsic developmental clock, orchestrating transitions between morphological attractors that ultimately give rise to outgrowth patterns required to build functional neural circuits.

### Periodic Branching Emerges from Sustained Entropic Equilibrium and Microstate Divergence

We next investigated whether a trajectory could be sustained along a boundary between two attractors over extended distances. Using the Monte Carlo simulation framework, we identified a highly complex, periodic branching phenotype (Figure 8A) where criticality is sustained. Initially, the trajectory is anterior. However, tracking the macrostate entropies along this trunk revealed that the system was not securely within a single basin of attraction. Instead, it continuously hovered near the phase boundary between anterior drive (macrostate 3) and ventral drive (Figure 8B). Periodically (Figure 8A, points 1-8), macrostate entropies equalized, and a physical bifurcation was triggered. At points 1 through 6, this equilibrium occurred between macrostate 3 and Macrostate 0. As the trajectory progressed farther into the gradient, the ventral drive shifted, resulting in an equilibrium between macrostates 3 and 6 at points 7 and 8. The diverging branches (Figure 8C, Trajectories 1-8) escaped the anterior attractor and fell fully into the ventral attractors (driven by macrostate 0 or 6). Trajectories 1 through 4 terminated prematurely before reaching the midline, due to the rising instability of the ventral macrostate 0 that matched the opposing dorsal drive of macrostate 5.

**Figure 8.**
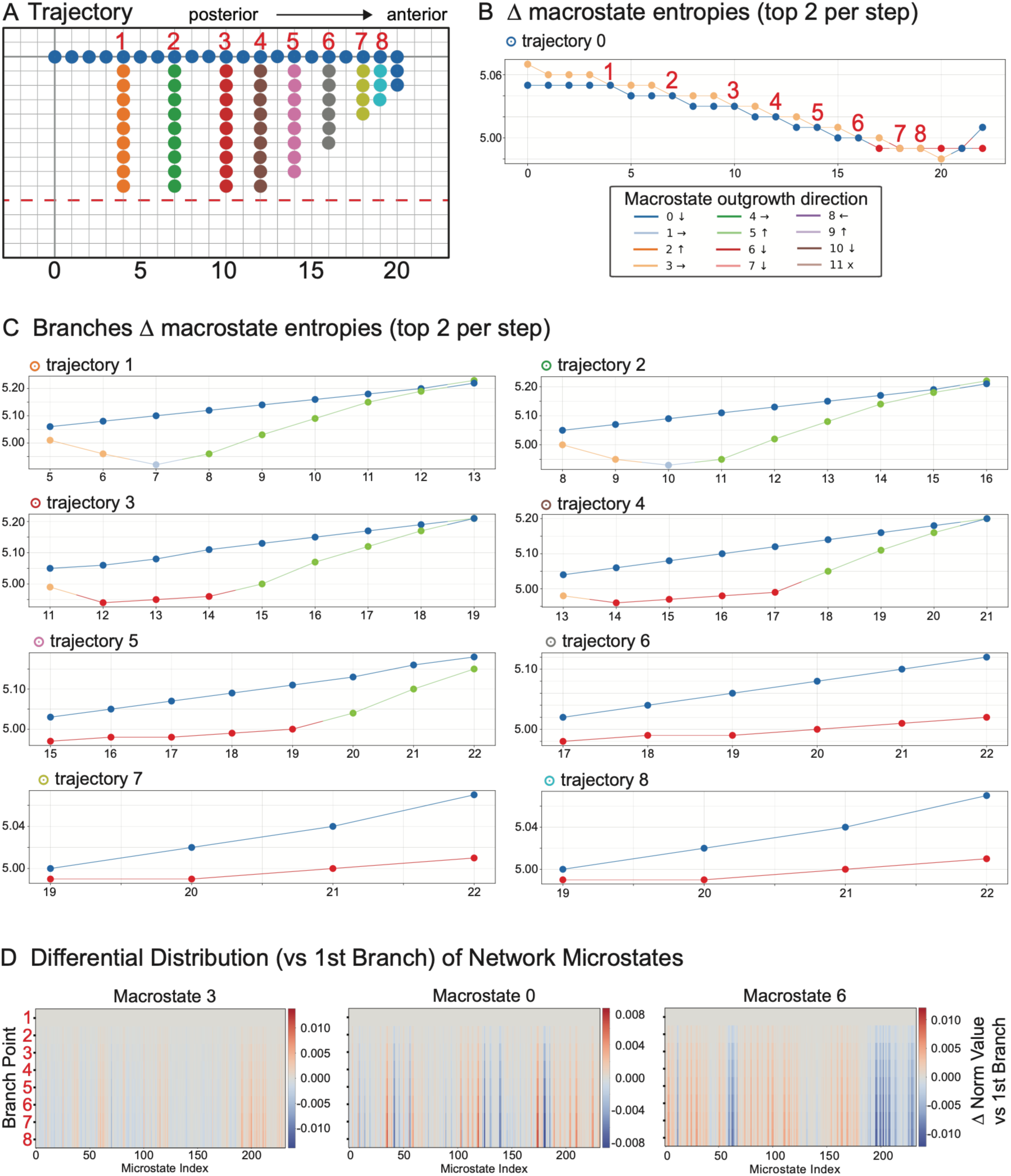
Sustained entropic equilibrium drives periodic branching and trajectory arrest. (A) A simulated 2D trajectory exhibiting a periodic branching phenotype. The primary anterior trunk (Trajectory 0) generates sequential ventral branches at spatial points 1 through 8. (B) Tracking of the top two macrostate entropies along the main trunk (Trajectory 0). An anterior drive maintains the trajectory (Macrostate 3) that periodically falls into perfect equilibrium with ventral drives (Macrostate 0 at points 1-6; Macrostate 6 at points 7-8), triggering the branching events. The overall macrostate entropy decreases along this primary axis. (C) Tracking of the top two macrostate entropies for the individual branches (Trajectories 1-8). Trajectories 1-4 experience premature arrest when the dominant ventral drive (Macrostate 0) reaches equilibrium with an opposing dorsal drive (Macrostate 5). The overall macrostate entropy increases as these branches progress. (D) Heatmaps displaying the differential distribution of the 232 network microstates comprising Macrostates 3, 0, and 6 at each branching point, calculated relative to the first branch (Point 1). The progressive increase in differential variance (red/blue intensity) demonstrates that the underlying network microstates continuously reconfigure to maintain macroscopic equilibrium as the growth cone navigates the changing extracellular gradient. Grid Orientation & Markers: In Panel A, the anterior direction is oriented to the right to align visually with the progression of the time axis. The intersection of the solid gray lines indicates the starting coordinate where all local extracellular cue values (UNC-6, SLT-1, EGL-20, WL) are equal to 1.0. The red dashed line denotes the ventral midline, corresponding to the spatial coordinates of maximum UNC-6 and minimum SLT-1 values. Values used for the plots are provided in Table 1.

Finally, we examined the network dynamics that enable the system to sustain this periodic branching behavior as it traverses a continuously changing extracellular environment. We analyzed the 232 network microstates underlying macrostates 3, 0, and 6 at each branching point. Heatmaps of microstate distributions reveal a progressive divergence across points 2–8 (Figure 8D). Although the anterior driver (macrostate 3) remains relatively stable, the internal composition of the ventral drivers (macrostates 0 and 6) shifts substantially over the trajectory. This computational result demonstrates the influence of network degeneracy. Moving through varying levels of extracellular cues might alter the macrostate with the highest macrostate entropy. However, the initial macroscopic entropic balance is preserved by the reconfiguration of microstate distributions. This allows a repeating pattern of branching to emerge.

## Discussion

Growth cones must remain both resilient and adaptable as they navigate extracellular environments filled with noisy and often conflicting cues. Our results suggest that the Guidance Signaling Network (GSN) operates at entropic phase boundaries. At these boundaries, the system can transition between macrostates that encode distinct directional programs. By moving along them, a growth cone can sustain straight-line trajectory while remaining poised to change direction. Even a modest input, such as a slight change in the level of an extracellular cue, can alter the dominant macrostate and change the pattern of outgrowth. Unlike models that impose explicit gradient-following rules, this framework determines the most likely trajectory based solely on the growth cone’s internal signaling state and local environmental cues. This principle enables the model to reproduce diverse in vivo patterning events, including turning, branching, bifurcations, bipolar outgrowths, regional specification, and choice-point navigation.

A central paradox in developmental neurobiology is how a migrating growth cone maintains a resilient, consistent trajectory while still retaining the plasticity to alter its path. Related to this is a contradiction between the complexity of neural circuit patterns and the apparently limited number of molecular guidance cues. Previously, we introduced the Statistically-Oriented Asymmetric Localization (SOAL) model, which frames navigation as a stochastic process driven by the probabilistic localization of force-producing machinery (25, 32, 37). Because the underlying molecular distribution is probabilistic, the growth cone’s overall trajectory can be mapped using the principles of statistical mechanics. Here, we have elevated the stochastic foundation into a comprehensive thermodynamic framework that is entropy-based. The macrostate entropy across 232 biologically viable network microstates was calculated at each spatial coordinate during a simulation of a growth cone transversing the extracellular environment. The results indicate that pathfinding is a continuous search for signaling instability. This allows the growth cone to maximize its available microstates and maintain plasticity precisely when a directional choice is needed.

When a specific morphological macrostate has low entropy, the GSN is confined to a narrow set of microstates, and the system is within a deep basin of attraction. Here, the growth cone is resistant to directional changes caused by genetic or environmental fluctuations because the actin cytoskeleton organization (the force-producing machinery) is controlled by a dominant signaling program that arises from the activity of only a few microstates. By contrast, the system sits near boundaries between attractors if it moves towards a macrostate with the highest entropy. This is a highly volatile state because here many microstates control signaling without anyone dominating. Small changes (chemical or mechanical) could trigger a new signaling program that rapidly reconfigures the actin active matter, leading to a directional turn.

A profound implication of our model is the growth cone’s ability to maintain a trajectory along an entropic phase boundary over extended spatial distances and to branch from the initial trajectory sequentially (Figure 8). Branching occurs when two competing macrostates reach equilibrium. As the growth cone migrates through shifting extracellular gradients, the specific distribution of the 232 network microstates comprising the dominant macrostates continuously reconfigures. The GSN is dynamically warping its internal state space to compensate for environmental changes. This allows the macroscopic entropic equilibrium to cycle along the initial trajectory. These results provide a computational demonstration of biological degeneracy, the ability of structurally distinct network configurations to yield identical macroscopic outputs.

While spatial gradients provide the environmental coordinates for pathfinding, development is strictly a function of time. In vivo, the precise sequencing of distinct migratory phases (e.g., cell body positioning followed by axon extension) is regulated by the temporal modulation of intrinsic factors. Our simulations modeling the temporal expression of the transcription factor zag-1 (Figure 7) captured this dynamic. The results demonstrate that progressively modulating the growth cone’s gene functions can shift the entropic landscape. The regulation of the transcription factor’s function acts as a developmental clock. Depending on the baseline and rate of zag-1 accumulation, the growth cone is pushed toward, or held back from, a critical phase transition. This illustrates that an intrinsic transcriptional program can interface with spatial environmental cues to sequentially unlock new basins of attraction during neurodevelopment.

While the entropic framework models the information-processing capacity of the Guidance Signaling Network (GSN), axon guidance ultimately requires the generation of mechanical work. From a biophysical perspective, the axonal growth cone functions as active matter. That is a driven, nonequilibrium assembly of actin filaments and microtubules, coupled to ATP-consuming motors and adhesion linkages that continuously convert chemical energy into force (38, 39). Studies suggest that the control of cytoskeletal organization provides the physical means by which signaling becomes motion (40–44). For example, actomyosin-driven flow and motor–clutch coupling appear to transduce guidance information into traction at point contacts, and locally biased microtubule dynamics can instruct turning decisions.

A defining property of active systems is the capacity for rapid symmetry breaking and non-equilibrium state transitions. The emergence of directed cortical flows in minimal actomyosin cortices, transitions between steady flows and periodic waves in confined active gels are examples of these cytoskeletal properties. We propose that entropic state changes arising from the GSN provide the informational triggers for the mechanical transitions in cytoskeletal active matter. When the GSN occupies a low-entropy basin (a stable attractor), a persistent directional polarity is maintained, perhaps through clutch engagement. Conversely, as the GSN approaches high entropy, cytoskeletal reorganization occurs, perhaps involving wave-like remodeling of actin and locally biased microtubule entry.

This perspective aligns with recent stochastic views of axon growth in which small spatial biases in intrinsically noisy actin networks integrate over time to yield net directional movement, rendering the growth cone sensitive to changes in cue-regulated probabilities rather than to deterministic instructions (37, 45). Ultimately, our findings suggest that the highly ordered wiring of the nervous system is paradoxically driven by stochastic, microscopic fluctuations in signaling networks poised at criticality.

In our framework, a growth cone’s trajectory is an emergent property of the collective guidance signaling network. Extracellular cues act nonlinearly to alter the probability distribution of network configurations. This in turn reshapes the entropy landscape that organizes the probabilities of alternative outgrowth behaviors. Canonical chemotropic behavior, defined as directed movement toward or away from an external chemical stimulus, emerges in the model when a high-entropy ridge aligns with a chemical gradient. Under these conditions, attraction and repulsion remain important descriptors of growth-cone behavior. At a molecular scale, conventional chemotropic models explain axon trajectories through mechanisms that generate attractive or repulsive responses to spatially distributed extracellular cues. The clear functional logic of the chemotropic framework has guided the discovery of many molecular mechanisms underlying axon guidance. For example, binding of the UNC-6 cue to the receptors UNC-40 and UNC-5 activates interacting signaling pathways. These interactions influence localized cytoskeletal organization and bias the direction of movement relative to the cue source. Our framework considers the same molecular components at a different level of organization. That is, how do the collective configurations of the molecular components give rise to macroscopic behavior? From this perspective, attraction and repulsion are context-and direction-dependent behavioral outcomes of the collective signaling network, rather than distinct guidance mechanisms with fixed relationships to particular molecules. Our framework explains how relatively few guidance cues can produce complex outgrowth patterns without requiring axonal trajectories to align consistently with chemical gradients or requiring cues to have fixed attractive or repulsive functions. Rather than converging towards a single dominant network configuration, persistent outgrowth follows high-entropy boundaries between competing behavioral macrostates. Network degeneracy along these boundaries allows multiple microscopic configurations to support the same directional-trajectory behavior. This also preserves access to the turning and branching states. We hypothesize that UNC-40 increases the diversity of accessible network configurations, whereas UNC-5 constrains the distribution enough to maintain a coherent directional behavior. Accordingly, the thermodynamic framework does not replace descriptions of directional behavior in terms of attraction and repulsion. Rather, it explains how these behaviors emerge under particular configurations of a dynamic signaling landscape.

## Limitations of the Computational Framework

Several limitations of the current modeling must be acknowledged. First, the simulation uses a simplified 2D spatial matrix. In contrast, in vivo neural environments are highly complex 3D tissue architectures in which both chemical gradients and mechanical topographies (e.g., substrate stiffness and physical extracellular matrix barriers) play roles. Second, the scalar values representing extracellular ligand gradients (UNC-6, SLT-1, EGL-20, WL) are approximations. The values are based on a scale where 1.0 represents the wild-type value for a cue during initial HSN axon outgrowth, and 0.0 represents the absence of the cue. The concentrations and slopes of these cues, and their relative proportions in vivo, remain technically difficult to quantify. (Although in some cases precise values might be needed to model a specific guidance, in other cases, the model’s qualitative behavior may be sufficient.) Finally, our 11-component GSN, while sufficient to capture macroscopic behavioral states, represents a highly abstracted fraction of the complete growth cone signaling network. Equally abstract is our current definition of the morphological macrostates themselves. In this study, the classification of complex in vivo outgrowth behaviors into 12 discrete, coarse-grained categories serves as a necessary but ‘rough’ approximation of the phenotypic space. Consequently, the quantitative association between a specific macrostate and a precise directional vector is not yet biologically rigorous. It is important to emphasize that the primary objective of this study was to establish the theoretical foundation for this approach, rather than developing a fully practical predictive model.

## Future Directions

The model presented in this study was trained exclusively on the genetic and morphological data of the *C. elegans* HSN neuron. The 11 components of the Guidance Signaling Network (GSN) used in this study are highly evolutionarily conserved molecules that govern axon guidance across organisms from nematodes to mammals. An interesting question is whether the non-linear associations learned by the Artificial Neural Network are unique to HSN or represent universally conserved rules of cellular navigation. Because our model calculates guidance decisions based on internal network volatility rather than rigid, hardcoded spatial rules, the theoretical framework of entropic steering is likely universal. However, in vivo, different neuronal classes navigate vastly different topographies; for instance, while the HSN navigates a gradient in which the SLT-1 (Slit) source is dorsal and UNC-6 (Netrin) is ventral, commissural neurons in many organisms navigate environments in which both Slit and Netrin are secreted simultaneously from the ventral midline. Future advancements will determine if a single, foundational neural network can be seamlessly adapted to distinct spatial arrangements. If not, the model may require a different approach, such as transfer learning, that recalibrates the baseline internal GSN weights for specific neuronal subtypes. Ultimately, scaling this approach from individual neurons to diverse neuronal classes could provide the computational foundation necessary to simulate the patterning of an entire connectome.

To transition this model from a theoretical framework to a precise, practical tool, future work must fundamentally improve the phenotypic measurements used to define the macrostates. The current model relied on coarse visual scoring of terminal phenotypes. Modeling could be improved by better quantification of cellular traits, enabling continuous rather than discrete mapping of phenotypic traits to the internal GSN microstates. This could be done by integrating high-throughput, automated morphometric analyses, such as computer vision-driven tracking of growth cone area, filopodial dynamics, and cytoskeletal turnover. The method used in this study provides an approach for systems biology. The approach could also be expanded by integrating real-time transcriptomic data to update the GSN’s intrinsic parameters. For example, to enable patient- or cell-type-specific simulations. Finally, expanding the GSN to include downstream cytoskeletal effectors could help bridge the gap between entropic signaling states and direct biophysical force generation.

The paradigm has broad implications for understanding other forms of complex cellular motility and has potential clinical applications. For example, cellular movements underlying neurodevelopmental wiring disorders, immune cell chemotaxis, and metastatic invasion could be similarly modeled. Disease states represent systemic failures of the cell to navigate appropriately localized entropic phase transitions. Current therapeutic strategies often rely on singular interventions, such as blocking a specific receptor. But this approach frequently has limited success due to the degeneracy and compensatory mechanisms encoded in genetic networks.

The computational framework established here provides a powerful in silico platform for uncovering these complexities. By mapping the entropy landscape of damaged tissue, this model could be deployed to run highly parallelized, combinatorial perturbation analyses. This could reveal precise combinations of network manipulations (e.g., targeted gene therapies or pharmacological interventions) required to shift the growth cone back to a critical state. Doing this could restore the cytoskeletal plasticity needed to navigate pathological environments and re-establish function.

## Experimentally Testable Predictions

Our theoretical framework conceptualizes axon guidance as a process driven by entropic phase transitions. The model simplifies complex molecular interactions that give rise to outgrowth patterns into coarse-grained thermodynamic properties. This paradigm can be rigorously tested, and potentially falsified, using biophysical and molecular techniques. For example: For example, we propose:

1. Non-Canonical Navigation in Engineered Gradients. Traditional chemotactic models suggest that growth cones navigate by descending or climbing gradients of attractant or repellent cues. Although it is recognized that the pathways mediating responses to these cues exhibit complex crosstalk, the assumption persists that cues encode a fixed, predictable direction. In contrast, our maximum-entropy steering algorithm predicts that growth cones actively seek states of high signaling volatility. Extracellular cues have no intrinsic directional properties. Experiment: Expose growth cones in a microfluidic device to different engineered, conflicting environments, such as orthogonal attractant and repellent fields. We predict that growth cones will not consistently follow a path determined by attractive and repulsive properties. Metric: Trajectories will instead conform to the calculated entropic phase boundary, migrating toward regions that explicitly maximize the macrostate entropy of the GSN.
2. Live-Imaging of Criticality and Mechanical Phase Transitions. We propose that shifts in the GSN’s entropic state serve as the informational triggers for mechanical phase transitions in cytoskeletal active matter. Experiment: Use multiple FRET biosensors to watch how key signaling molecules fluctuate, while concurrently observing cytoskeletal changes in live cells. Falsifiable Metric: During straight-line migration (low-entropy basins), cytoskeletal flow will exhibit a persistent steady state. However, spikes in cytoskeletal volatility that occur during spontaneous symmetry breaking or branching will not be random. They will be immediately preceded by a massive, quantifiable spike in signaling variance (critical fluctuations) that maps to the spatial coordinates of an entropic phase boundary.
3. Single-Cell Validation of Network Degeneracy. Our framework illustrates that growth cones maintain prolonged trajectories along phase boundaries by continuously reconfiguring their underlying microstates. This network degeneracy implies that the growth cone can rely on various combinations of receptors and signaling molecules to produce the same morphological behavior at different points along a gradient. Experiment: Isolate individual growth cones executing the exact same morphological behavior (e.g., a specific turn) and perform rapid single-cell multiomic profiling. Falsifiable Metric: Traditional models predict that identical growth cones will have the same internal states when executing the same turns. In contrast, our model predicts degeneracy, in which individual growth cones exhibit statistically distinct combinatorial microstates despite sharing a macrostate. Some guidance receptors may appear to act redundantly, at least under the conditions where the guidance is observed. We predict that selectively inhibiting such a receptor could cause abrupt, localized guidance defects. These failures will occur at spatial coordinates where the growth cone’s dynamically warped state space depends heavily on a microstate that requires that receptor’s activity.
4. Manipulation of Entropic Crossovers. Our simulations demonstrate that the modulation of intrinsic factors (such as the transcription factor *zag-1*) functions as a developmental clock that can predictably shift the entropic landscape. Experiment: Use optogenetics or other rapidly inducible systems to either prematurely increase the concentration of these transcription factors *in vivo*, or artificially “clamp” specific GSN nodes into fixed activity states. Falsifiable Metric: A change in the system’s macrostate entropy can lead to unusual morphological behaviors, such as premature branching or abrupt directional changes. In contrast, clamping a node limits the accessible microstates, thereby reducing the system’s entropy. This would enforce a straight-line trajectory, even in environments where new patterns might be expected to emerge.

## Materials and Methods

### Dataset Construction and Pre-processing

The MLP training dataset was from in vivo genetic studies that quantified axon outgrowth in C. elegans. The input feature space consisted of 11 binary variables representing the genetic state of specific guidance cues and receptors (*egl-20, mig-1, sax-3, slt-1, unc-40, unc-5, unc-53, unc-6, wl, wr, zag-1*). The target variable corresponded to one of the 12 morphological outgrowth patterns (the 11 observed topologies plus an isotropic “null” state) that corresponded to the geneotype (Figure 2B). Resampling with replacement was used to equalize class frequencies. The balanced dataset was partitioned into training (85%) and validation (15%) subsets.

Because neural network models can produce misleading predictions when input signals are absent, we included an isotropic ‘null’ macrostate in the training dataset. This macrostate represents an unbiased random walk paired with a synthetic ‘null’ genotype, with the directional cue thevalues set to 0. By incorporating this into the training regimen, the MLP was trained to recognize the specific pattern left by missing information. Consequently, if a simulated growth cone moves to a spatial coordinate where extracellular cue values are all zero, the model defaults to the isotropic macrostate. This simulates a stalled growth cone in which filopodia extend and retract randomly due to internal noise.

### Iterative Model Generation

We built multi-layer perceptrons (MLPs) using the Keras Functional API (TensorFlow). Each model comprised an 11-unit input layer, a 200-unit ReLU-activated dense layer, a Dropout layer (rate = 0.6), and a 12-unit Softmax output layer. To comprehensively probe the gene-network topology, we trained >1,000 model instances. Each model was initialized with random weights and trained using the Adam optimizer (learning rate=0.001) and Sparse Categorical Crossentropy loss for 80 epochs (batch size=7).

### Ensemble Calibration and Selection

To calibrate the simulation engine, models were rigorously evaluated against the unseen test set. We selected a final cohort of five models based on a dual criterion: (1) individual predictive accuracy exceeding 95%, and (2) complementary coverage, where the selected models exhibited non-overlapping failure modes (i.e., deficiencies in one model were covered by the predictive strengths of another).

These five models were loaded into a custom ensemble architecture. The ensemble arranges the five base models in parallel, sharing a common input. The probability vectors (outputs) of the five constituent models are combined via an averaging layer (layers.Average()), yielding a final consensus probability distribution. This ensemble model was saved and used as the decision-making engine for all subsequent simulations.

### Model Validation via Random Walk Simulations

We validated the MLP model by comparing its predictions to in vivo experimental data in the probabilistic random walk model. Simulations were conducted to calculate the average displacement of a theoretical growth cone navigating a 2D plane.

The probabilities of outgrowth determined the movement vectors for the random walk in four cardinal directions: anterior (−x), posterior (+x), ventral (−y), and dorsal (+y). Two parallel sets of simulations were executed. The first set utilized the directional outgrowth probabilities directly observed from in vivo genetic mutant experiments. The second set used the probabilities predicted by the trained MLP model for the combinatorial binary input vectors corresponding to genotypes of the mutants analyzed in vivo.

To translate the MLP’s macrostate predictions into directional movements needed for the random walk simulation, the SoftMax probabilities for each state were weighted by the in vivo observed probabilities for each direction. The resulting directional sums were then normalized to sum to 1 to serve as transition probabilities for the random walk. The resulting spatial coordinates (average Δx and Δy, with standard deviations) for each genetic combination were plotted to visually and quantitatively evaluate the concordance between the experimental and computationally derived directional biases (Figure 3). Python scripts utilizing NumPy and Pandas were used to calculate the walk trajectories, and Matplotlib was used for visualization.

### Entropic Steering Algorithm

The steering mechanism that uses the MLP model to explore a spatial environment was implemented in Python. The Entropic Steering Algorithm simulates axon pathfinding as a discrete-time walk on a 2D grid. The grid represents the extracellular environment, with coordinate axes corresponding to gradients of guidance cues (e.g., UNC-6, SLT-1, EGL-20, WL).

1. State Evaluation and Biologically Constrained Combinatorial Dropout At each simulation step, the algorithm samples the local extracellular cue concentrations at the current spatial coordinate (x, y). These environmental scalar values are concatenated with the base intrinsic cellular gene states to form a composite input vector. To assess the robustness of the GSN at this specific spatial location, the algorithm performs a combinatorial dropout analysis. A perturbation ensemble (M) is generated by systematically substituting wild-type intrinsic gene states (1.0) with null states (0.0). Because we want a biologically relevant phase space, the perturbation ensemble was restricted to only single, double, and triple null-mutant combinations. Combinations of four or more null mutations are typically lethal and do not contribute to viable macrostates. This generates a localized “mutational neighborhood” consisting of exactly 232 distinct network microstates (M = 232) surrounding the base input vector at every spatial coordinate.
2. Calculation of Macrostate Entropy As conceptually detailed in the Results, macrostate entropy (*H_k_*) was used to quantify network degeneracy. Computationally, this was executed at each spatial coordinate (*x*, *y*) by iteratively querying the trained MLP. A custom script (Python using TensorFlow) generated an input array for the 232 defined perturbation microstates based on local gradient concentrations. The MLP outputted a probability distribution across the 12 morphological macrostates for each perturbation. To calculate *P_norm_*(*m*|*k*), the raw probability for a specific macrostate was divided by the sum of probabilities for that macrostate across all 232 microstates. For the Shannon entropy calculation, to prevent undefined values when *P_norm_* = 0, these zero-probability microstates were filtered out and excluded from the calculation prior to the natural logarithm transformation.
3. Directional Mapping and Execution The algorithm uses these calculated entropies as the sole steering metric. The macrostate exhibiting the maximum Macrostate Entropy (H_max_) is selected as the dominant biological drive for that time step. The winning macrostate k_max_ is then mapped to a physical cardinal direction (Anterior, Posterior, Dorsal, Ventral, or Stop). Because biological responses to extracellular cues are context-dependent, this mapping is dynamically adjusted based on the growth cone’s spatial domain (e.g., distinct mapping rules are applied depending on whether the coordinate is pre-or post-midline crossing). Once a direction is determined, the algorithm updates the (x, y) coordinates of the simulated growth cone by one step. This sample-evaluate-move cycle repeats until the trajectory reaches the matrix grid boundary or a designated maximum step limit. The result is a pathway generated from purely stochastic local interactions.

To capture the physical constraints of axonal extension, the algorithm imposes a self-avoiding walk (SAW) during trajectory generation. A spatial memory matrix records all previously visited (x, y) coordinates for a given trajectory. If the maximally entropic movement vector directs the growth cone into a previously occupied space, the step is rejected, and the growth cone stalls. This SAW constraint computationally prevents local minima and infinite oscillatory loops. Biologically, it reflects contact inhibition and the physical reality that a growth cone cannot retract into its own stabilized, microtubule-rich axon shaft.

### Monte Carlo Simulation of Intrinsic State Tuning and Choice Point Navigation

To explore whether the entropic steering algorithm can generate the morphological diverse patterns observed in nature, Monte Carlo simulations were used to sample the intrinsic cellular gene-parameter space stochastically. The spatial matrix representing the extracellular environment (UNC-6, SLT-1, EGL-20, and WL gradients) was held constant across all simulations. Intrinsic gene values were randomly assigned from a continuous uniform distribution, and trajectories were predicted (Figure 5, Panels A-P).

To model localized state changes at anatomical boundaries, a choice point navigation algorithm was implemented (Figure 5, Panels Q-T). Simulations were initialized with all intrinsic parameters set to 1.0, corresponding to the wild-type state that drives a simulated growth cone to ventrally toward the midline. The midline choice point is at the spatial coordinate with the maximum UNC-6 value and the minimum SLT-1 value. The algorithm pauses at the midline and the wild-type intrinsic gene values are replaced with values stochastically generated via Monte Carlo sampling. This simulates an environmental factor at the midline altering the cell’s signaling network. The entropic steering algorithm then resumes, using these updated intrinsic parameters to calculate the modified microstate entropies and determine subsequent movement vectors. Trajectories crossing the midline utilized an expanded spatial matrix encompassing both the right and left sides of the simulated anatomical axis.

### Visualization of Entropic Dynamics and Phase Transitions

To correlate spatial trajectory changes with internal network dynamics, the Macrostate Entropy (H_k_) for all 12 morphological macrostates was recorded at every discrete step of the simulation. To provide a comprehensive visualization of the system’s volatility, the continuous entropy values for all 12 states were plotted against simulation time steps (trajectory coordinates). To isolate and identify entropic phase transitions (crossover events), a sorting algorithm was applied at each time step to extract the two macrostates with the highest entropy values. These “Top 2” values were plotted longitudinally. An entropic phase transition was defined mathematically as the specific time step at which the rank order of the maximum-entropy state and the second-highest entropy state inverted. These crossover time steps were subsequently mapped back to the 2D spatial trajectory to confirm their alignment with physical directional changes.

### Simulation of Temporal Intrinsic State Dynamics and Branch Tracking

To model the temporal accumulation of transcription factors during development, the entropic steering algorithm was modified to accept dynamically updating intrinsic parameters. For time-series simulations evaluating the transcription factor zag-1 (z1), the baseline intrinsic values for all other cellular genes were held constant at 1.0. At each sequential time step (equivalent to one spatial movement execution), the scalar value representing z1 was incremented according to predefined continuous gradients (e.g., ranging from 0.025 to 1.3875).

Following each temporal update of the z1 parameter, the macrostate entropies for all 12 macrostates were recalculated. The simulation employed a branch-tracking queue system. If the algorithm identified multiple macrostates with maximal equivalent entropy, the physical trajectory was split into multiple trajectories. Unique Trajectory IDs were assigned to the diverging paths, and the simulation iteratively computed the subsequent spatial and entropic steps for each independent branch.

### Differential Network Microstate Analysis and Heatmap Visualization

To investigate the internal network composition underlying complex periodic behaviors, we tracked the distribution of network microstates at specific spatial coordinates during trajectory bifurcation. At each branching point, the normalized output probabilities (microstate logits) for the 232 distinct network microstates comprising the dominant macrostates were extracted from the MLP.

To examine how the internal state space changed throughout the periodic trajectory, we carried out a differential analysis. The microstate distribution at the initial branching event (Point 1) was used as a reference. We then subtracted this baseline from the normalized 232-microstate vectors at each later branching point (Points 2–8) to obtain the differential distribution (Δ Norm Value). The differential heatmaps show that network configurations become increasing divergent, even as the system produced comparable macroscopic morphological outcomes.

### Model Evaluation

Performance was quantified using Sparse Categorical Accuracy. The ensemble strategy was validated by demonstrating that the averaged output of the five complementary models successfully classified problematic inputs that individual networks could not. All analysis was performed in Python using Pandas, Scikit-learn, and TensorFlow/Keras.

## Data availability

The software archive is available at Zenodo: 10.5281/zenodo.22923634

Computational Framework for a Thermodynamic Theory of Axon Guidance 2026-09-23

## Acknowledgments

We thank The Office of Advanced Research Computing (OARC) at Rutgers, The State University of New Jersey, for providing access to the Amarel cluster and associated research computing resources, which have contributed to the results reported here. URL: https://it.rutgers.edu/oarc. We thank David J. Foran, PhD, and the Biomedical Informatics Shared Resource at Rutgers Cancer Institute for providing access to dedicated compute nodes to conduct these studies.

## Funding

The data presented in Figure 1 were collected with support from the National Institutes of Health (NIH) National Institute of Neurological Disorders and Stroke, grant numbers NS-033156 and NS-061805.

Portions of the text were refined with the assistance of AI language tools (Microsoft Copilot and Google Gemimi). Python script was generated with the assistance of PyCharm AI Assistant (JetBrains) and Google Gemini. All scientific content, data interpretation, and final text were reviewed, validated, and finalized by the author.

## Author Contributions

W.G.W designed and performed research, analyzed data, and wrote the paper.

